# Exquisite Sensitivity to Dual BRG1/BRM ATPase Inhibitors Reveals Broad SWI/SNF Dependencies in Acute Myeloid Leukemia

**DOI:** 10.1101/2021.04.23.441171

**Authors:** Florencia Rago, Lindsey Ulkus Rodrigues, Megan Bonney, Kathleen Sprouffske, Esther Kurth, GiNell Elliott, Jessi Ambrose, Peter Aspesi, Justin Oborski, Julie T. Chen, E. Robert McDonald, Felipa A Mapa, David A. Ruddy, Audrey Kauffmann, Tinya Abrams, Hyo-eun C. Bhang, Zainab Jagani

**Affiliations:** Novartis Institutes for Biomedical Research, Cambridge, MA, USA; Novartis Institutes for Biomedical Research, Basel, Switzerland

**Keywords:** SWI/SNF, BRG1/SMARCA4, BRM/SMARCA2, AML

## Abstract

Various subunits of mammalian SWI/SNF chromatin remodeling complexes display loss-of-function mutations characteristic of tumor suppressors in different cancers, but an additional role for SWI/SNF supporting cell survival in distinct cancer contexts is emerging. In particular, genetic dependence on the catalytic subunit BRG1/SMARCA4 has been observed in acute myeloid leukemia (AML), yet the feasibility of direct therapeutic targeting of SWI/SNF catalytic activity in leukemia remains unknown. Here, we evaluated the activity of dual BRG1/BRM ATPase inhibitors across a genetically diverse panel of cancer cell lines and observed that hematopoietic cancer cell lines were among the most sensitive compared to other lineages. This result was striking in comparison to data from pooled short hairpin RNA screens, which showed that only a subset of leukemia cell lines display sensitivity to BRG1 knockdown. We demonstrate that combined genetic knockdown of BRG1 and BRM is required to recapitulate the effects of dual inhibitors, suggesting that SWI/SNF dependency in human leukemia extends beyond a predominantly BRG1-driven mechanism. Through gene expression and chromatin accessibility studies, we show that the dual inhibitors act at genomic loci associated with oncogenic transcription factors, and observe a downregulation of leukemic pathway genes including *MYC*, a well-established target of BRG1 activity in AML. Overall, small molecule inhibition of BRG1/BRM induced common transcriptional responses across leukemia models resulting in a spectrum of cellular phenotypes. Our studies reveal the breadth of SWI/SNF dependency and support targeting SWI/SNF catalytic function as a potential therapeutic strategy in AML.

## Introduction

The mammalian SWI/SNF complexes (also known as the BAF complexes) are ATP-dependent chromatin remodelers that facilitate transcriptional regulation of gene expression through repositioning of nucleosomes at promoters and enhancers. SWI/SNF is typically comprised of 12-15 subunits, which assemble into distinct functional configurations depending on cell lineage and context. Whole exome sequencing studies have detected SWI/SNF subunit mutations in ∼20% of cancers (1,2), and functional genetic screens have identified dependencies on paralogous SWI/SNF subunits in these mutated contexts. These include BRM (*SMARCA2*) dependence in *BRG1* (*SMARCA4*)-deficient and ARID1B (*BAF250B*) dependence in *AR1D1A* (*BAF250A*)-deficient tumor models (3-7). In contrast, dependence on SWI/SNF subunits BRG1 and BRD9 in the absence of any known SWI/SNF mutations has been observed in hematopoietic and lymphoid cancer models (8-10). In the hematopoietic lineage, BRG1 plays both an essential role as a critical regulator of hematopoiesis during development (11,12) and in leukemogenesis (9,13,14). In leukemia models, BRG1-dependent remodeling of the *MYC* enhancer region has been shown to regulate expression of *MYC*, suggesting a potential mechanism for BRG1 dependency in AML (9). These findings suggest that the mechanisms underlying SWI/SNF dependency vary with genetic context.

In this work, we utilize dual BRG1/BRM catalytic inhibitors (15,16) to further investigate SWI/SNF dependency in hematopoietic malignancies. We find that the majority of hematopoietic cancer cell lines evaluated are exquisitely sensitive to SWI/SNF inhibition. Further, we elucidate through genetic studies that the sensitivity to the inhibitors is not driven solely through BRG1 dependence, but also relies on concomitant loss of BRM function. Using a focused panel of AML cell lines, we demonstrate that BRG1/BRM inhibitors modulate common leukemic transcriptional programs across cell lines, inducing a broad range of phenotypic outcomes including differentiation and apoptosis. Finally, the observation that BRG1/BRM inhibitor treatment slows tumor growth in an AML xenograft mouse model suggests therapeutic potential for SWI/SNF inhibition, and sheds light on important pharmacokinetic/pharmacodynamic considerations for future drug candidates.

## Materials and Methods

### Cell lines and culturing

All cell lines were SNP verified and mycoplasma tested. See Supplementary Table S1 for culture conditions for each cell line.

### Cell line engineering

The control, BRG1, and BRM shRNA sequences used were described previously (3,17). shRNAs were cloned into a pLKO-Tet-On vector system (3) containing a U6 promoter and either puromycin (puro) or neomycin (neo) selection markers. The MYC overexpression construct, based on a previously described expression vector (18), was synthesized (GeneArt) and cloned into a lentiviral vector driven by an EF1alpha/HTLV promoter with a blasticidin selection marker. Lentiviral production was described previously (3). Lentiviral transduction was performed by plating 400,000 cells/well in 24-well deep round-bottom plates in 1 ml growth media containing 8 µg/ml polybrene. Virus (500 µl) was added and cells were spun with lentivirus at 2250 rpm for 90 minutes at room temperature. Following centrifugation, an additional 1 ml of media was added. Cells were allowed to recover for 3 days before selection with puro (1 µg/ml), G418 (0.5 mg/ml), or blasticidin (10 µg/ml). For combined knockdown of BRG1 and BRM, MOLM-13 cells were first infected with shBRG1-puro lentiviruses; following recovery from selection, the cells were then infected with shBRM-neo lentiviruses and selected with G418. shRNA expression was induced with 100 ng/mL doxycycline.

### Cell line profiling

Cells were plated at 250 cells per well in 1536-well plates in duplicate. The next day, cells were treated with BRM011, BRM014 or BRM017 (11-point dose titration, 3.16-fold dilutions). After 3 days, cell viability was measured using Cell Titer-Glo Luminescent Cell Viability Assay (CTG, Promega) on a luminescence plate reader. Viability was calculated relative to DMSO-treated cells (100% viability) and MG132-treated cells (3.33 mM, -100% viability). Amax and Crossing point (concentration at which viability curve crosses 50%) were calculated from the resulting viability curves.

### PBMC isolation and activation

Whole blood was obtained from donors who have provided full informed consent for research purposes via IRB-approved Informed Consent Form and protocol. Peripheral blood mononuclear cells (PBMC) were isolated by centrifugation at 1800xg for 25 min at room temperature in Heparin-coated CPT tubes (BD Biosciences). Samples were diluted with 1x PBS and centrifuged at 1500 rpm for 5 minutes. PBMC pellet was subsequently resuspended in PBS. To activate cells, CD3/CD28 magnetic beads (ThermoFisher Scientific) were added to PBMCs (cell/bead ratio = 8:1).

### 5 day proliferation assays

For 5-day proliferation assays, cells were plated in 40 µl of growth media in 384-well plates (Corning). See Supplementary Table S1 for plating density for each cell line. Cells were treated with BRM011, BRM014, or BRM017 (11-point, 3-fold serial dilutions) in triplicate using an Echo550 (Labcyte). Viability was assessed on Day 0 and Day 5 using CTG according to manufacturer’s instructions. Luminescence was measured using a PheraStar FSX instrument (BMG Labtech). For 24-hour proliferation assays, cells were plated at a density of 2×10^4 cells/well. For PBMC proliferation assays, activated PBMCs were plated at 10,000 cells per well in a 384-well plate and treated/processed as above. PBMCs were stimulated with CD3/CD28 Dynabeads to induce proliferation before treatment. Viability was measured at 0, 3 and 5 days. Inhibition (Inh) values were calculated by normalizing to the untreated wells from the Day 5 timepoint for the given cell line. Growth inhibition values were calculated as described previously (19). Normalized data were fit using the three parameters nonlinear regression function in GraphPad Prism. AAC50 values are reported as concentrations of compound where curve fit crosses 0.5.

### RT-qPCR

Cells (2×10^4) were plated in phenol-free RPMI (Life Technologies) with 10% FBS in 384-well plates and treated with varying doses of BRM011, BRM014, or BRM017 in triplicate using an Echo550. After 24 hours, cells were lysed by adding 3x lysis buffer (10 mM Tris-HCl, 1.5% Igepal, 150 mM NaCl, 1 U/µl RNasin (Promega)). Reverse transcription and quantitative PCR (RT-qPCR) was performed using 0.25 µl of lysate and Taqman Fast Virus 1-step Master Mix (Applied Biosystems) on a CFX384 thermal cycler (Bio-Rad). Probes used for gene expression analyses are listed in Supplementary Table S2.

### RNA sequencing (RNA-seq)

Cells (1⨯10^6 in 1.5 ml growth media) were seeded in triplicate in 24-well deep round-bottom plates and treated with BRM011 or equal volume DMSO. BRM011 doses were chosen to induce >90% *MYC* down-regulation (30 nM HL-60, MOLM-13, HEL 92.1.7, Kasumi-1, CMK-86, OCI-AML-3; 200 nM SKM-1; 300 nM THP-1). After 24 hours, cells were harvested by centrifugation at 4°C followed by cell lysis/RNA extraction using RNEasy plus mini kit (Qiagen) according to manufacturer’s instructions. Samples were eluted in 30 µl of water. RNA was quantified and assessed for purity using the RNA ScreenTape on the Agilent 4200 TapeStation System. Sequencing and analysis of RNA-seq samples were performed as described previously ((17,17), see Supplementary Materials and Methods).

### Assay for Transposase-Accessible Chromatin with high-throughput sequencing (ATAC-seq)

THP-1 cells (2×10^6 in 3 ml growth media) were seeded in triplicate in 24-well deep round-bottom plates and treated with DMSO or BRM011 (300 nM). After 24-hour treatment, cells were processed, sequenced and analyzed as described previously ((17), see Supplementary Materials and Methods).

### Western blot

Standard techniques were used for protein extraction and immunoblotting (see Supplementary Materials and Methods). Antibodies used were: MYC (Abcam ab39688), BRG1 (Abcam EPR3912), BRM (Cell Signaling D9E8B), Vinculin (Sigma V9131), PARP (Cell Signaling 9542), GAPDH (Millipore MAB374), Immun-Star Goat Anti-Mouse-HRP conjugate (Bio-Rad), and Immun-Star Goat Anti-Rabbit-HRP conjugate (Bio-Rad).

### Fluorescence activated cell sorting (FACS)

Cells (1-1.5×10^6) were incubated with FACS sample buffer (DPBS, 0.5% FBS) and human FC Block (Invitrogen) for 20 minutes on ice followed by incubation at 4°C for 30 minutes with the following - antibodies: anti-CD34-FITC, anti-CD38-BV421, anti-CD45RA-BV650, anti-CD135-PE, anti-CD33-PE/Cy7, and anti-CD11b-APC (all from BioLegend). Cells were then washed twice with FACS sample buffer. Annexin V and propidium iodide staining was performed with AlexaFluor 488 Annexin V/Dead Cell Apoptosis Kit (ThermoFisher V13241) per manufacturer’s instructions. All FACS analyses were performed on a BD LSRFortessa. Data was analyzed using FlowJo (version 10).

#### *In vivo* efficacy study

Mice were maintained and handled in accordance with the Novartis Institutes for BioMedical Research (NIBR) Institutional Animal Care and Use Committee (IACUC) and all studies were approved by the NIBR IACUC. Female SCID-beige mice (Charles River) were acclimated in NIBR animal facility (12 hour light/dark cycle) with ad libitum access to food and water for at least 3 days before manipulation. Mice (6–8 weeks old) were inoculated subcutaneously in the right dorsal axillary region with the MV-4-11 cell line (20 10^6^ cells in 50% Matrigel). Tumor volumes and body weights were monitored twice per week and the general health condition of mice was monitored daily. Tumor volume was determined by measurement with calipers and calculated using a modified ellipsoid formula, where tumor volume (TV) (mm3) = [((l w2) 3.14159))/6], where l is the longest axis of the tumor and w is perpendicular to l. When average tumor volume reached approximately 200 mm^3^, animals were randomly assigned to receive daily dosing of either vehicle or BRM014 20 mg/kg. Compound treatments began 11 days post MV4-11 cell implantation. Tumor samples of the BRM014 treated group were collected for PD analysis post 7 and 14 days of daily dosing (4 and 24 hours post the last dose). Tumor samples of the vehicle group were also collected after 7 and 14 days of daily dosing (4 hours post final dose).

### RT-qPCR from *in vivo* samples

RNA extraction, cDNA synthesis and RT-qPCR was performed as described previously ((17), see Supplementary Materials and Methods). Probes used are described in Supplementary Table S2.

### Processing of *in vivo* samples for Western blot

T-PER (Invitrogen) lysis buffer + 1x HALT protease/phosphatase inhibitor cocktail (Thermo Fisher Scientific) was added to tumors samples, which were then homogenized using Lysing Matrix D beads using a Precellys 24 homogenizer. Samples were then further homogenized by passing through Qiashredder columns (Qiagen) and probed for MYC expression by Western blot as described above.

## Results

### Cell line profiling reveals sensitivity of hematopoietic cancer cell lines to dual BRM/BRG1 ATPase inhibitors

Previous work demonstrated that a subset of hematopoietic cancer cell lines are dependent on BRG1 (9), however, analysis of a large panel of cell lines screened with pooled short hairpin RNAs (shRNAs) did not demonstrate a uniform lineage wide dependency on either SWI/SNF catalytic subunit, *BRG1/SMARCA4* or *BRM/SMARCA2* (Figure 1A, (20)). Interestingly, a survey of several core and accessory SWI/SNF subunits (*SMARCE1, ARID2* and *SMARCC1*) showed a more consistent pattern of dependency across hematopoietic cell lines (Figure S1A). Together, these data suggest a dependency on the SWI/SNF complexes as a whole, rather than on a particular complex member. To further test this hypothesis, we took advantage of well characterized dual BRG1/BRM ATPase inhibitors, BRM011 and BRM014 (15,16) to evaluate growth inhibition across 100 cancer cell lines from various cancer primary sites (21). After 3 days of treatment, we observed a profound sensitivity across all the hematopoietic lines tested (N=27) to either BRM011 or BRM014 (Figure 1B, S1B, Table S3). In contrast, the cell inactive analog BRM017 (15) was significantly less active in this panel (Figure S1B, Table S3). To further evaluate the sensitivity observed in hematopoietic cancer cell lines, we treated a smaller AML-focused panel of cell lines (N=16) with BRM011, BRM014 and BRM017 for five days and measured the effect on viability. The small panel comprised a variety of different genetic backgrounds (Table S4). We observed a wide range of sensitivities across this panel (Table S5), and classified the lines into three categories based on their absolute AC50 (AAC50) of growth inhibition: highly responsive, moderately responsive and weakly responsive (Figure 1C, S1C). Interestingly, cell lines across all categories varied in depth of response; 9/16 cell lines underwent a growth arrest, with Amax ≥ 0 (e.g. MOLM-13, Figure 1C), while 7/16 lines showed Amax ≤ 0 (e.g. Kasumi-1, Figure 1C), suggestive of cell death.

**Figure 1:**
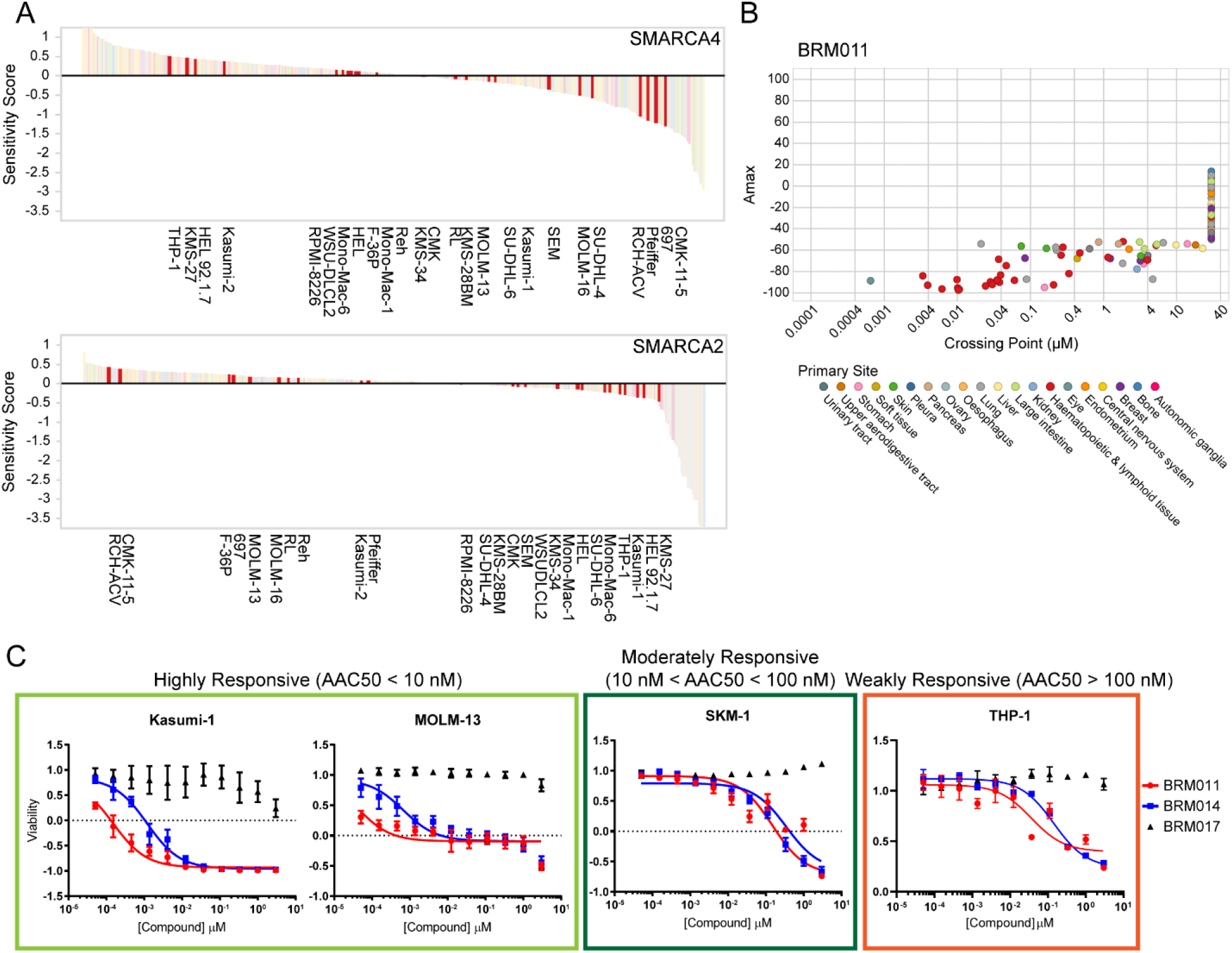
Hematopoietic cancer cell lines show a dependence on SWI/SNF by both genetic and chemical perturbation. (A) Data from DRIVE pooled shRNA screens (20) plotted as sensitivity score for indicated genes across all cell lines. Hematopoietic cancer cell lines are highlighted in red and labeled. (B) Amax plotted versus crossing point (point at which curve crosses y = 50%) across cell lines treated with BRM011 for 3 days. Values were calculated by curve fitting with DMSO-treated cells as 100% and MG132-treated cells as -100% viability. Cell lines are colored by primary site as indicated. Cell line identities together with Amax and crossing point values are listed in Supplementary Table S3. (C) 12-point dose response curves for representative hematopoietic cells treated with BRM011, BRM014, and BRM017 (N=3 per treatment, error bars shown as s.d.). Cell lines were categorized as follows: highly responsive AAC_50_ < 10 nM (light green), moderately responsive 10 nM < AAC50 < 100 nM (dark green), weakly responsive AAC50 > 100 nM (orange).

Finally, to understand whether the observed effects on proliferation were specific to the hematopoietic lineage, we tested normal human PBMCs with the same compounds. While we did observe growth inhibition in the normal PBMCs, these cells fell into the moderately responsive category (Figure S1D, Table S5). This suggests that while normal hematopoietic cells are also dependent on SWI/SNF, the transformed state in a subset of hematopoietic cancers can impart a profound sensitivity to SWI/SNF inhibition.

### Dual knockdown of BRG1 and BRM phenocopies effects of BRG1/BRM inhibitors

Surprisingly, several cell lines including MOLM-13, KMS-34, Reh and HEL 92.1.7, which did not show genetic dependence on either BRG1 or BRM individually in the pooled shRNA screen (Figure 1A), were sensitive to the dual inhibitors in proliferation assays. First, we examined *BRG1* and *BRM* mRNA expression under basal culture conditions across the cell line panel (Figure 2A-B). We observed a range of expression levels that did not appear to predict dependency. To further investigate the discrepancy between genetic and chemical perturbation, we chose to engineer AML cell line MOLM-13 with *BRG1* and *BRM* targeting shRNAs. MOLM-13 is not responsive to knockdown of either BRG1 or BRM alone in pooled shRNA screens (Figure 1A), but shows sub-nanomolar AAC50 to SWI/SNF inhibition (Table S5). MOLM-13 was engineered to express doxycycline inducible shRNAs against *BRG1* and *BRM*, either alone or in combination. Suppression of gene expression was confirmed by RT-qPCR (Figure 2C-D). As expected, knockdown of BRG1 or BRM alone did not affect proliferation of MOLM-13 cells over the course of a 6-day growth assay; however, dual knockdown of BRM and BRG1 resulted in significant reduction in cell proliferation (Figure 2E), similar to what was observed with the dual BRG1/BRM inhibitor treatment. Importantly, these data demonstrate that depletion of multiple SWI/SNF subunits concurrently may be required to unmask dependency in AML models, further supporting the importance of the family of SWI/SNF complexes in AML.

**Figure 2:**
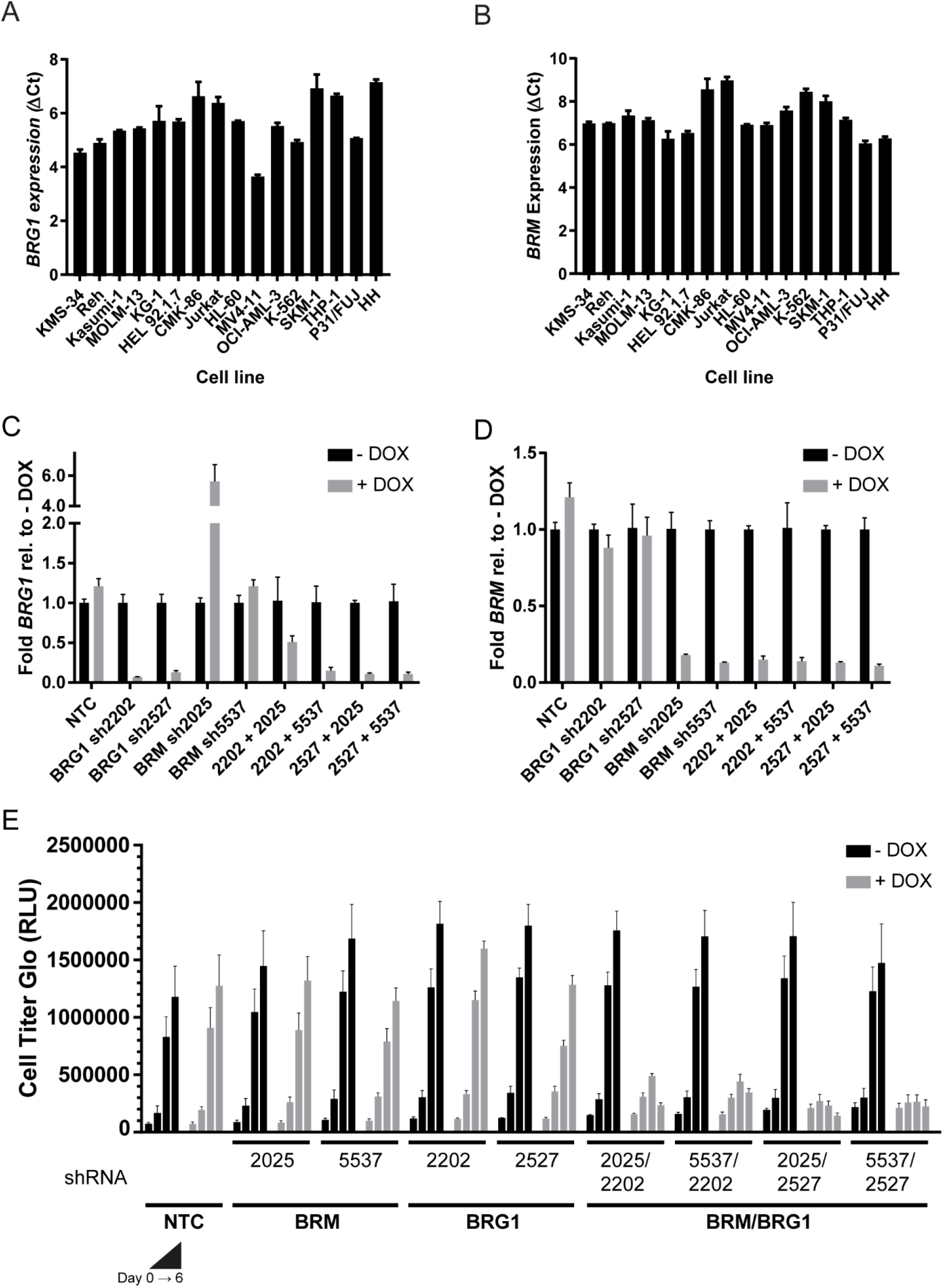
Dual knockdown of BRG1 and BRM inhibits growth of AML cells that are not affected by knockdown of BRG1 or BRM alone. (A-B) Basal BRG1 (A) or BRM (B) gene expression in hematopoietic cell line panel. Ct values were normalized to β-actin and plotted as delta Ct (N=3 biological replicates per condition, error bar shown as s.d.). (C-D) qRT-PCR analysis of BRG1 (C) or BRM (D) gene expression in MOLM-13 cells containing dox-inducible shRNAs against BRG1 and/or BRM or a non-targeting shRNA control (NTC) harvested 48 hours post-dox treatment. Gene expression was normalized to β-actin and plotted as fold change (2^-ddCt) relative to no dox condition for each cell line (N=3 biological replicates per condition, error bars shown as s.d.). (E) Proliferation assay time course of the engineered MOLM-13 cells lines measured by Cell Titer Glo (N=3 per condition, error bars shown as s.d.). Time points (from left to right) are Day 0, 2, 4, 6 post-dox treatment.

### Response to dual BRM/BRG1 inhibitors extends beyond regulation of MYC expression

Previous studies suggested that the direct regulation of *MYC* gene expression by BRG1 underlies the dependency observed in various hematopoietic malignancies (9). We tested this by treating our cell line panel with low dose (0.004 µM) or high dose (3 µM) BRM011, BRM014 or BRM017 for 24 hours and measured the effect on *MYC* mRNA transcript levels. Of note, BRG1/BRM inhibitors did not decrease *BRG1* gene expression (Figure S2A) at this time point. We found that across all the cell lines tested, *MYC* expression was downregulated upon treatment with either BRM011 or BRM014, but not BRM017 (Figure 3A). Only the cutaneous T cell lymphoma line HH showed no change in *MYC* expression, consistent with the lack of growth inhibition in this cell line (Figures S1C). We further tested *MYC* regulation in response to SWI/SNF inhibition by treating cells in dose response with BRM011. As seen in the viability assays, we observed varied depth of response across the cell lines, although the range of AC50s was more narrow (Figure S2B, Table S6). Interestingly, there was no significant correlation between growth inhibition and basal *MYC expression*, AC50, or depth of *MYC* modulation (Figure S2C).

**Figure 3:**
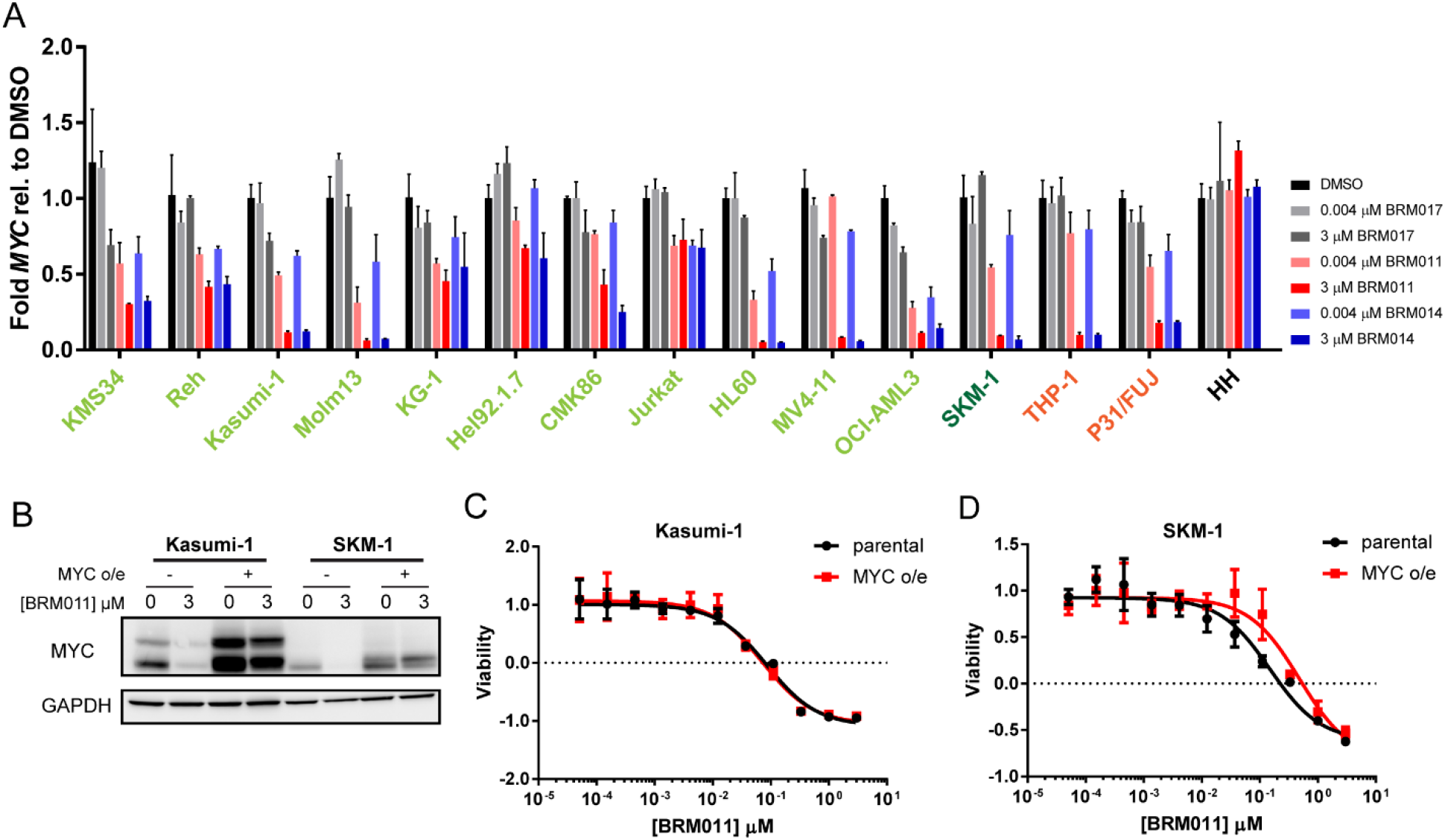
SWI/SNF inhibition decreases *MYC* gene expression across hematopoietic cancer cell lines. (A) RT-qPCR analysis of *MYC* expression in hematopoietic cancer cell lines treated with DMSO, 0.004 µM or 3 µM BRM011, BRM014 and BRM017 for 24 hours. *MYC* expression was normalized to *ACTB* and plotted as fold change relative to DMSO treated (2^-ddCt) (N=3, error bars shown as s.d.). (B) Immunoblot of lysates from Kasumi-1 and SKM-1 parental or MYC overexpression cell lines +/- 3 µM BRM011 for 24 hours probed with anti-MYC or anti-GAPDH antibodies. (C-D) 5-day proliferation assays for Kasumi-1 (E) or SKM-1 (F) parental or MYC overexpression cell lines treated with BRM011 at indicated doses (N=3, error bars shown as s.d.).

To further probe the disconnect between growth inhibition and *MYC* modulation, we overexpressed MYC in Kasumi-1 (highly responsive) and SKM-1 (moderately responsive) cells (Figure 3B). Exogenous overexpression rescued the MYC protein downregulation observed after compound treatment (Figure 3B), and led to a slight growth advantage for the MYC overexpressing cells (Figure S3A-B). However, this overexpression did not rescue the viability effect in 5-day growth assays when treated with BRM011 or BRM014 (Figure 3C-D, S3C-F). These data suggest that while *MYC* expression is responsive to SWI/SNF modulation as previously described, neither *MYC* expression, nor *MYC* modulation is an accurate predictor of inhibitor sensitivity in the models examined.

### AML cell lines can undergo both differentiation and apoptosis upon SWI/SNF inhibition

Next, we investigated the phenotypes of various human hematopoietic cancer cell lines upon SWI/SNF inhibition. Previous reports in a mouse MLL-AF9 AML model indicated that depletion of BRG1 by gene knockdown led to differentiation of the cancer cells (9,13). However, it was unknown if human hematopoietic cancer cell lines would undergo differentiation or apoptosis when treated with a BRG1/BRM inhibitor, and our viability data suggested that the phenotype may be cell line dependent (Figure 1C, S1C). To determine whether the cell lines were undergoing apoptosis, we first probed a panel of cell lines for PARP (Poly (ADP-ribose) polymerase) cleavage. After treatment with 3 µM BRM011 for 24 hours, we observed an increase in cleaved PARP in only a subset of lines (Figure 4A). Surprisingly we did not detect cleaved PARP in all lines that appeared to undergo cell death in viability assays, so we additionally probed for apoptosis by Annexin V/propidium iodide (PI) staining. Cells were treated for 48 hours with DMSO or BRM011. JQ1, a BET inhibitor known to induce cell death in AML cell lines (8), was used as a positive control. We observed an increase in Annexin V single-positive cells and Annexin V/PI double-positive cells in AML models upon BRM011 addition (Figure 4B, S4A-H). Only Reh and Jurkat, ALL and T cell leukemia lines, respectively, did not show an increase in Annexin V staining with BRM011 or JQ1 (Figure S4I, J), consistent with PARP cleavage data. These data demonstrate that SWI/SNF inhibition can promote apoptosis in AML cell lines.

**Figure 4:**
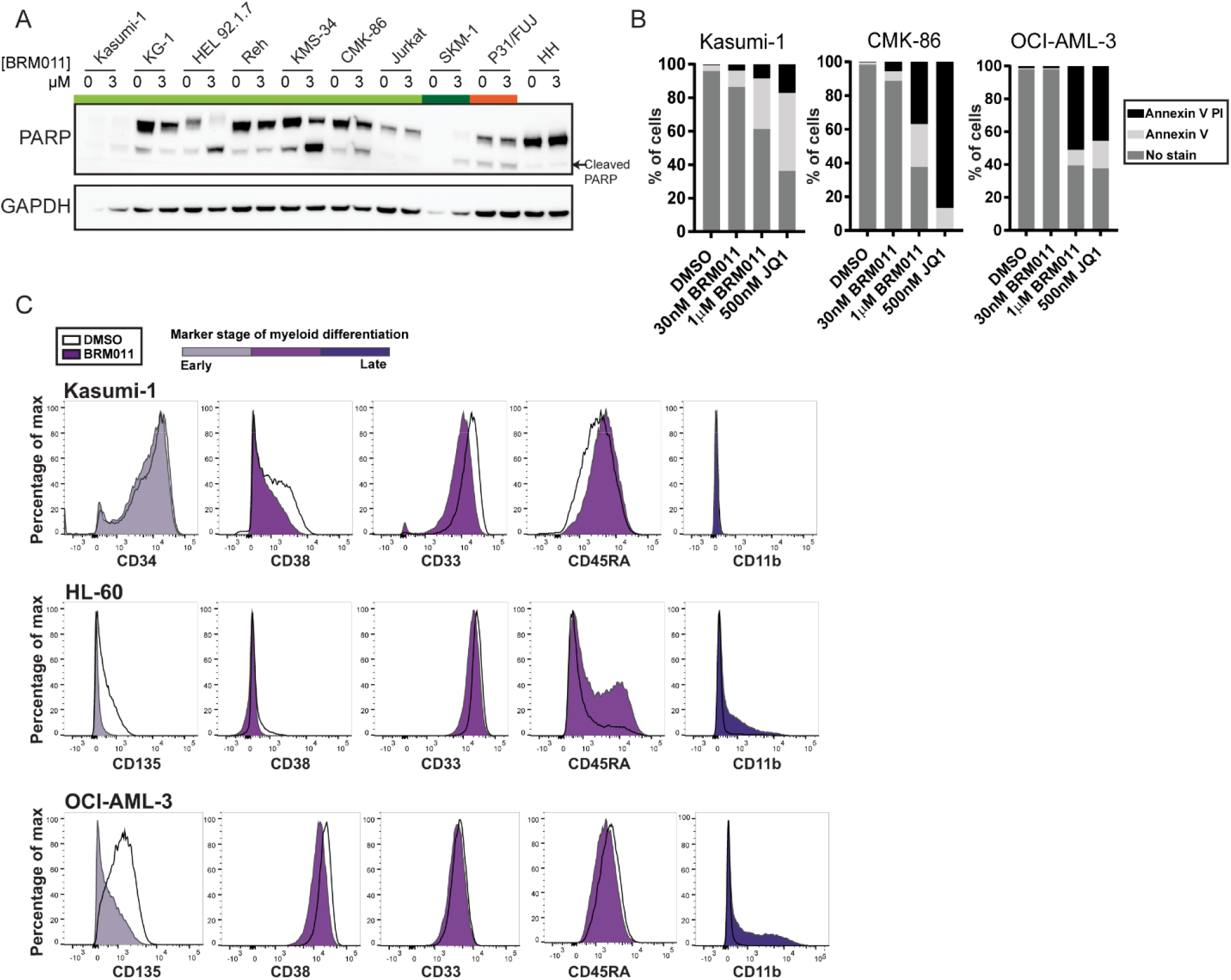
SWI/SNF inhibition by BRM/BRG1 ATPase inhibitors induces apoptosis and differentiation. (A) Immunoblot of hematopoietic cell line lysates treated with DMSO or 3 µM BRM011 for 24 hours and probed with anti-PARPor anti-GAPDH. (B) Percentage of cells unstained, Annexin V stained or Annexin V and propidium iodide (PI) stained as measured by flow cytometry after 48 hours in DMSO or indicated doses of BRM011. JQ1 (500 nM, 48 h) was used as positive control. (C) Flow cytometry analysis of cells treated with DMSO (white histogram) or 30 nM BRM011 (purple histograms) for 72 hours and stained with a panel of cell surface markers representing various stages of hematopoietic differentiation: CD34, CD135, CD38, CD33, CD45RA and CD11b.

We further probed the response to inhibition of SWI/SNF activity by testing modulation of cell surface proteins associated with different stages of hematopoietic lineage development. Lineage marker expression was assessed in a panel of cell lines by flow cytometry after 72 hours of treatment with DMSO or BRM011 (initial survey of marker expression annotated in Supplementary Table S7). Markers of immaturity, such as CD34 and CD38, tended to decrease in the presence of BRM011 (Figure 4C, S4L-V). In cell lines with measurable basal expression, CD135 levels strongly decreased (Figure 4C, S4L, O-Q, T, V). CD135, encoded by the *FLT3* gene, is a commonly mutated receptor tyrosine kinase that promotes leukemic proliferation, and its decrease upon BRM/BRG1 ATPase inhibition could potentially contribute to the growth inhibition observed in AML lines such as MOLM-13 that are *FLT3* mutated. CD45RA, which can serve as a marker of granulocyte monocyte progenitors (GMPs), increased within a subset of BRM011-treated Kasumi-1 and HL-60 cells (Figure 4C) (22). This may signal differentiation into the granulocyte monocyte lineage from an earlier stage in hematopoietic development following BRG1/BRM inhibition. However, this was not a universal response as other cell lines displayed decreased CD45RA following BRM011 treatment (Figure S4L, M, O, Q, R, V). Intriguingly, a cell line-specific response was observed for the monocytic marker CD11b, often used as a marker of differentiation into the macrophage lineage. Cell lines with a deeper decrease in viability with compound treatment, such as Kasumi-1 (Figure 1D), were less likely to increase CD11b-positive cells (Figure 4C, S4L-Q, U). Cell lines with a stasis-like phenotype, such as HL-60 and OCI-AML-3, increased CD11b levels following BRM011 addition compared to DMSO (Figure 4C, S4R-T, V). Overall, BRG1/BRM inhibition visibly induced differentiation in only a subset of lines, and showed a more cell line-specific response than the Annexin V apoptosis data.

### Genome wide analysis reveals common oncogenic pathways altered by BRM011 treatment in AML cell lines

Thus far, we have demonstrated that SWI/SNF perturbation affects cell viability and differentiation of AML cell lines and that the phenotypic effects observed in response to BRG1/BRM inhibition are likely not driven through modulation of *MYC* gene expression alone. To investigate how SWI/SNF inhibition more broadly affects gene transcription, we performed RNA sequencing (RNA-Seq) on 9 AML cell lines treated with DMSO or BRM011 for 24 hours. Consistent with our RT-qPCR data, we observed that *MYC* as well as *MYC* transcriptional target genes were significantly downregulated in Kasumi-1, OCI-AML-3, CMK-86, HL-60, SKM-1 and THP-1 cells (Figure S5A, red dots). However, *MYC* and its downstream targets were not significantly altered in MOLM-13, HEL or P31/FUJ cells treated with BRM011. To determine what pathways were altered in response to BRG1/BRM inhibition in AML, we next analyzed the significantly up- and downregulated genes in each cell line (abs(logFC) ≥ 0.5, p < 0.01) and further analyzed those genes using the Molecular Signatures Database (MSigDB) Hallmark gene sets. As expected, the MYC oncogenic pathway was significantly downregulated in 4 out of 9 cell lines, including Kasumi-1, SKM-1 and THP-1 (Figure 5A-D, Table S8). Other pathways significantly downregulated in response to BRG1/BRM inhibition across multiple cell lines included MTORC1, IL2/STAT5, KRAS, and EMT pathways (Figure 5A-D, Table S8). Additionally, hematopoietic metabolism was significantly upregulated in 5 out of 9 cell lines, indicating that BRG1/BRM inhibition has hematopoietic lineage-specific effects (Figure 5D). To further investigate consequences of BRM011 on hematopoietic-specific differentiation, we probed a 17-gene AML leukemic stem cell (LSC) gene set (23) and observed downregulation of this signature in a subset of cell lines (Figure S5B). In general, there were a number of commonly altered pathways in response to BRG1/BRM inhibition, but cell lines did not always group together according to pathway altered. For instance, Interferon gamma response (red dots, Figure 5A-C) was downregulated in 4 cell lines including SKM-1 and THP-1 cells, but not Kasumi-1. This observation lends support to the findings from the proliferation and differentiation assays described above; AML cell lines are exquisitely sensitive to BRG1/BRM dual inhibitors but the resulting phenotypic changes are diverse, illustrating the heterogeneity of AML.

**Figure 5:**
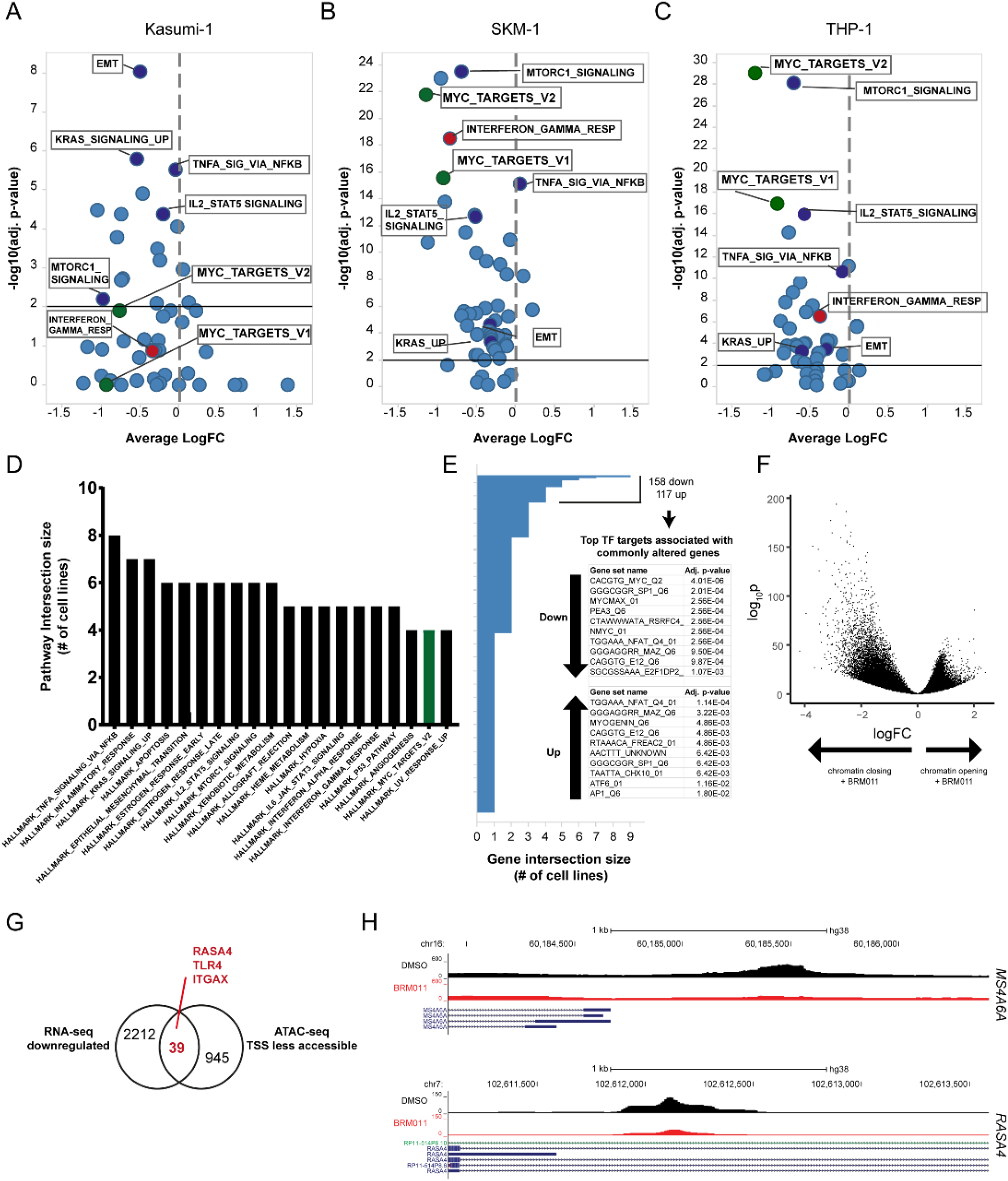
Common pathways are regulated downstream of SWI/SNF inhibition. (A-C) Volcano plots showing Hallmark pathways enriched for differentially expressed genes in BRM011-treated versus DMSO-treated Kasumi-1 (A), SKM-1 (B), and THP-1 (C) cells (24 hour treatment) measured by RNA-Seq. Average logFC is plotted versus -log10[adjusted p-value] for each pathway; p<0.01 was considered significant (black solid line). Hallmark_MYC_TARGETS_gene sets are highlighted in green. Examples of pathways commonly altered are highlighted in dark blue (common to all 3 cell lines) or red (shared between SKM-1 and THP-1 only). (D) Bar graph illustrating the top commonly altered Hallmark pathways across all 9 cell lines with number of cell lines that are altered for a particular pathway plotted on y-axis. Identities of cell lines in each group can be found in Table S8. (E) Bar graph indicating commonly altered genes across cell lines. Topmost significantly enriched transcription factor motifs are shown for the 158 genes downregulated and the 117 genes upregulated in at least 4 cell lines (GSEA analysis). (F) Volcano plot from ATAC-seq analysis showing more and less accessible chromatin regions in THP-1 cells upon treatment with 300 nM BRM011 for 24 hours. (G) Comparison of RNA-seq downregulated genes and ATAC-seq TSS less accessible genes. (H) Examples of less accessible regions at *MSRA6A* and *RAS4A* promoters following BRM011 treatment.

The SWI/SNF complex interacts with both general and hematopoietic-specific transcription factors including EKLF, RUNX1, PU.1, and CEBPα (9,24,25). To explore this further, we identified 158 downregulated and 117 upregulated genes that were commonly altered in 4 or more cell lines in our RNA-Seq dataset and performed GSEA analysis for transcription factor targets. Top transcription factor-binding motifs significantly enriched in both the up- and down-regulated gene sets included MYC, SP1, NFAT and MAZ (Figure 5E). We then evaluated chromatin accessibility changes in response to BRG1/BRM inhibition by performing ATAC-Seq analysis on THP-1 cells treated with BRM011 under the same conditions as RNA-Seq (300 nM, 24 hours) and conducting a genome-wide analysis to compare accessibility between samples. BRM011 treatment induced both chromatin closing and opening across the genome (8.3% of peaks less accessible compared to 5.3% more accessible upon BRM011 treatment) (Figure 5F). To identify transcription factor-binding motifs enriched in the chromatin regions that underwent changes in accessibility upon compound treatment, we ran HOMER with the default set of motifs and identified transcription factor binding motifs that were significantly enriched against a matched background (230 motifs in the less accessible regions and 160 in the more accessible regions; p < 0.01) (Table S9-10). Among the significantly enriched motifs, MYC, SP1, NFAT and MAZ scored in the analyses of both more and less accessible chromatin regions, consistent with the RNA-Seq enrichment analysis. Additionally, we observed enrichment of the known heme-specific SWI/SNF interactors, EKLF, RUNX1, PU.1, and CEBP further supporting the role of SWI/SNF in hematopoietic-transcription factor driven expression patterns. Finally, direct comparison of our RNA-seq and ATAC-seq data for THP-1, focusing on the genomic regions around the transcription start site (TSS), identified 39 genes with both decreased accessibility at their promoter region and transcriptional downregulation (Figure 5G) including *RASA4, MS4A6A, ITGAX*, and *TLR4* (Figure 5H, S5A, C). We similarly identified 7 genes that displayed increased promoter accessibility and transcriptional upregulation (Figure S5A, D-E). Collectively, these data demonstrate that BRG1/BRM inhibition alters chromatin accessibility and transcription at genes associated with general oncogenic and leukemia-specific transcriptional programs across leukemia models of varying genetic backgrounds.

### BRM014 treatment decreases the growth of human AML tumor xenografts in vivo

To evaluate whether consequences of chemical perturbation of the SWI/SNF complex *in vitro* translated *in vivo*, we tested the effects of BRM014 treatment on the growth of MV4-11 AML tumor xenografts. Since BRM014 is better suited to *in vivo* dosing, we confirmed *MYC* modulation upon BRM014 treatment first *in vitro* (Figure S6A) and saw similar repression of mRNA expression as we observed with BRM011 (Figure S2B). MV4-11 cells were implanted as subcutaneous xenografts and once tumors reached 200 mm^3^, mice were dosed once daily with 20mg/kg BRM014. Treatment continued for 14 days, and tumors were collected at several time points after the last dose (Figure 6A). We observed a significant reduction in tumor volume (T/C 51.7%) (Figure 6B) with no effect of treatment on mouse body weight (Figure S6B). Furthermore, robust decrease in *MYC* mRNA and protein expression was observed both during treatment and after study termination (Figure 6C-D, S6C-D). However, the dosing schedule was not sufficient to sustain MYC repression as MYC levels returned to untreated levels within 24 hours of dosing. These results are consistent with an *in vitro* washout experiment, in which *MYC* levels began to rebound at 7 hours post-washout, with *MYC* expression at basal levels by 17 hours post-BRM014 dose (Figure S6E). Collectively, these data demonstrate that sustained repression of SWI/SNF may be required for tumor regression, which will require further exploration of dose scheduling with future drug candidates. Overall, these studies have important implications for the potential of therapeutic SWI/SNF targeting in AML, expanding the opportunities to overcome the limitations of current treatments for this devastating disease.

**Figure 6:**
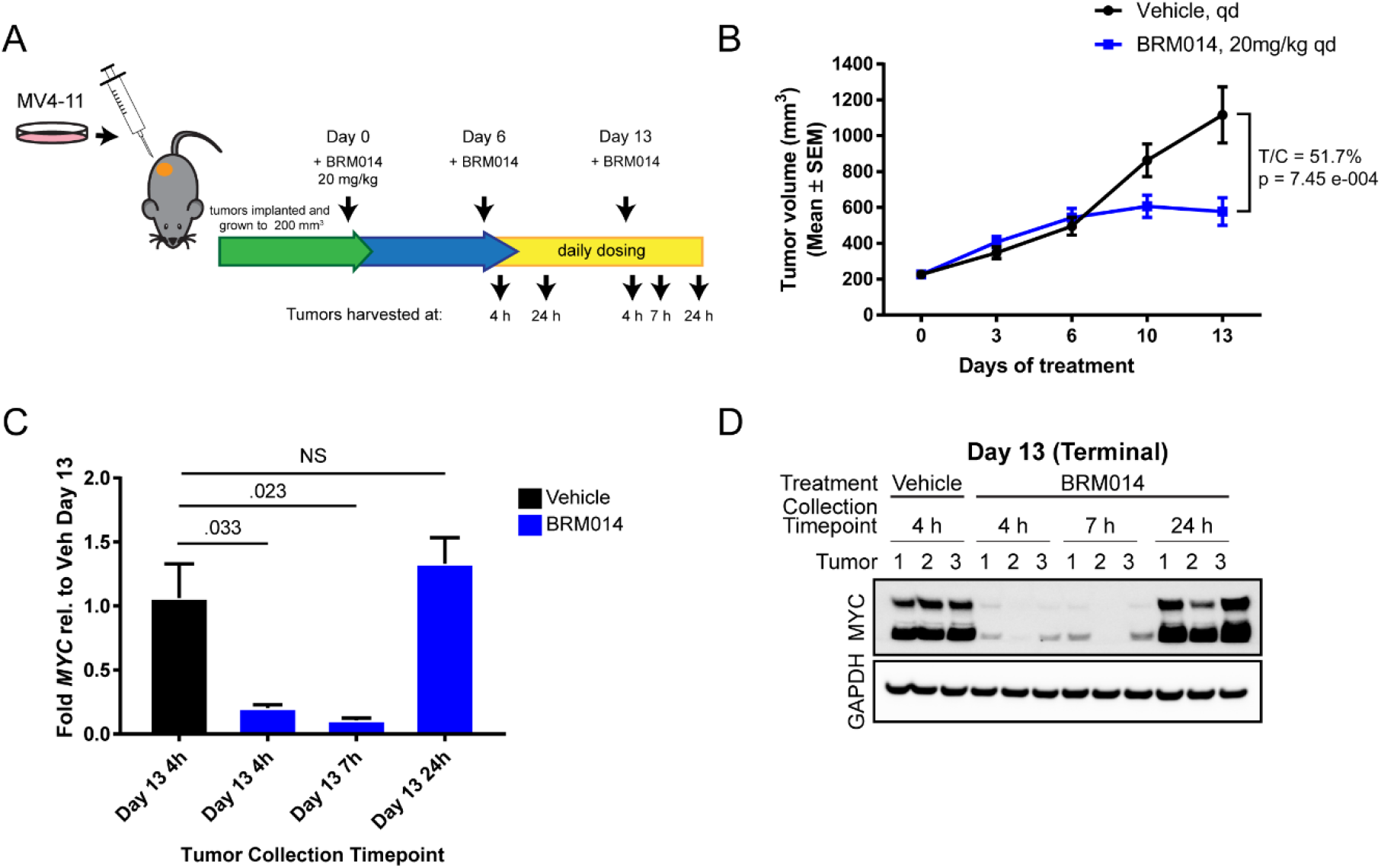
*In vivo* efficacy of BRM014 in AML xenograft model. (A) Dosing scheme for *in vivo* treatment of MV-4-11 with BRM014. (B) Tumor volume measurements (mm^3^) following treatment with vehicle or BRM014 (20mg/kg) for 2 weeks. (N=7 mice in vehicle group and 9 mice in treatment group, error bars shown as s.e.m). (C) Fold change in *MYC* expression across tumors treated with vehicle or BRM014 measured by RT-qPCR. MYC expression was normalized to *ACTB* and plotted as average fold change relative to vehicle-treated tumors (error bars shown as s.e.m., Student’s two tailed t-test, p-value indicated). (D) Immunoblots of MYC and PARP levels in mice treated with vehicle or BRM014 for labeled periods of time. Each column represents tumor from a unique mouse. GAPDH was used as a loading control.

## Discussion

Epigenetic de-regulation (i.e. recurrent mutations in TET2, DNMT3, IDH1/2, EZH2, ASXL-1) is a common feature of AML and other leukemias. In addition to the commonly mutated epigenetic regulators, AML models have been shown to be quite sensitive to perturbation of the SWI/SNF complex by both knockdown approaches (9,20) and chemical inhibition (10). Here, using dual BRG1/BRM ATPase inhibitors to interrogate a genetically diverse panel of AML and other hematopoietic cancer models we demonstrate that SWI/SNF dependency is broader than previously described. Through broad cell line profiling, we found that the hematopoietic lineage is significantly more sensitive than cancers from other primary sites to SWI/SNF inhibition. This phenotype is dependent on the activity of our compounds against both BRG1 and BRM subunits, resulting in more penetrant suppression of SWI/SNF catalytic activity. Interestingly, conditional dual knockout of Brg1 and Brm in the adult mouse hematopoietic compartment had no adverse effects on stem cell renewal or hematopoietic lineage differentiation (26), suggesting a potential sparing of normal hematopoietic tissue upon treatment with SWI/SNF inhibitors. Ultimately, the potential for a therapeutic window with SWI/SNF targeting will have to be rigorously tested with drug candidate small molecule inhibitors.

In this study we find that contrary to previous reports suggesting that SWI/SNF perturbation results in differentiation, the spectrum of phenotypes is much more varied. Some lines undergo surface marker expression changes indicative of differentiation to the monocyte/macrophage lineage, while others display more mild phenotypic changes and yet others undergo apoptosis. Additionally, interrogation of a panel of AML cell lines across our study uncovered the complexities of transcriptional response to SWI/SNF inhibitor treatment, beyond *MYC* modulation alone. We observed profound changes in transcriptional programs that affected a broad spectrum of oncogenic pathways (e.g. KRAS, MTORC1, EMT) as well as leukemia-specific transcriptional programs (e.g. PU-1). Remarkably, many oncogenic genes and pathways altered by BRM011 in hematopoietic models, such as KRAS, EMT and TNFα/NFKβ pathways, were similarly affected by this compound in BRG1-deficient lung cancer models (15), suggesting a consistent mode of action for BRG1/BRM ATPase inhibitors across cell lineages. Collectively, our characterization of the SWI/SNF dependency using the BRM/BRG1 ATPase inhibitors illustrates the variability in hematopoietic cancer cell responses to alterations in epigenetic regulators, likely driven by genetic and phenotypic heterogeneity of these hematopoietic malignancies.

Finally, we show successful growth inhibition of an AML xenograft model *in vivo* with BRM/BRG1 inhibitor treatment, demonstrating the efficacy of SWI/SNF inactivation in a tumor model. Importantly, we find that treatment with our inhibitor is not sufficient to cause regression in this model; however, this is likely due to insufficient repression of SWI/SNF-dependent gene expression, as evidenced by *MYC* expression rebound, and could be remediated if SWI/SNF complex inhibition is durably maintained over the course of the treatment. Furthermore, combination treatment with other epigenetic modulators that impair tumor growth in hematopoietic models may improve *in vivo* efficacy, although these treatments would likely need to be tailored to the specific genetic lesions found in each tumor. Apoptosis-inducing agents that target the BCL2 family or MDM2/p53 pathway have shown promise in AML models (27), and could also be evaluated in combination with BRG1/BRM inhibitors. Overall, this work demonstrates the potential for SWI/SNF inhibition as a treatment for AML and supports further research into compounds that can more durably inhibit complex activity, and combinations able to deepen the effect of single agent activity.

## Supporting information

Supplementary Figured and Tables

## Acknowledgements

The authors thank Jiang Zhu, Vesselina Cooke and Josh Korn for helpful discussions and Ye Wang, Pu Zhang, and Frank Chung for technical assistance.

