## Supplementary Materials and Methods for "Exquisite Sensitivity to Dual BRG1/BRM ATPase Inhibitors Reveals Broad SWI/SNF Dependencies in Acute Myeloid Leukemia"

**RNA sequencing**

200 ng of high purity RNA (RNA Integrity Number 7.0 or greater) was used as input to the Illumina TruSeq Stranded mRNA Library Prep Kit, High Throughput (#RS-122-2103), and the sample libraries were generated per manufacturer’s specifications on the Hamilton STAR robotics platform. The PCR amplified RNA-Seq library products were then quantified using the Advanced Analytical Fragment Analyzer Standard Sensitivity NGS Fragment Analysis Kit (#DNF-473). The samples were diluted to 10 nM in Qiagen Elution Buffer (Qiagen material #1014609), denatured, and loaded at a range of 2.5-4.0 pM on an Illumina cBOT using the HiSeq® 4000 PE Cluster Kit (#PE-410-1001). The RNA-Seq libraries were sequenced on a HiSeq® 4000 at 75 base pair paired end with 8-base pair dual indexes using the HiSeq® 4000 SBS Kit, 150 cycles (#FC-410-1002). The sequence intensity files were generated on instrument using the Illumina Real Time Analysis software. The resulting intensity files were demultiplexed with the bcl2fastq2 software and aligned to the human transcriptome using PISCES version 2018.04.01.

**Differential Expression and Pathway Enrichment Analysis**

Differential expression was determined using limma from Bioconductor (PMID: 25605792). Genes were called differentially expressed if they had an average Log2 expression >= 1 (reported as AveExpr in limma), an adjusted p-value <= 0.01, and an absolute Log2 fold-change of 0.5 relative to DMSO control. Gene set enrichment analysis for differentially expressed genes was performed using the hypergeometric test with FDR-adjusted p-values for pathway sets downloaded from MSigDB (PMID:16199517; Hallmark).

**ATAC-sequencing**

ATAC-seq was performed using a modified Omni-ATAC-seq protocol (1). Briefly, cells were treated with DNAse I (Worthington) for 30 min and then 1x10^5 cells per sample were isolated for subsequent steps. Nuclei isolation, tagmentation, and purification were performed as described in (1). Transposed DNA fragments were amplified in a 50 ul reaction containing using Nextera Index kit primers (Illumina), Nextera Sample prep kit PCR primer cocktail (Illumina) and NEBNext High-Fidelity 2x PCR mastermix (New England Biolabs). Number of amplification cycles was determined individually for each sample by qPCR using SYBR green and calculated as number of PCR amplification cycles required to one-third maximum fluorescent intensity. All libraries were amplified with <11 cycles to ensure sufficient library complexity. After amplification, libraries were purified using Agencourt AMPure XP beads (Beckman Coulter) at a sample to beads ratio of 1:1.2 and recovered in 12 µl of Buffer EB (Qiagen). Final product quality and concentration was determined using the High Sensitivity D5000 DNA Tape on the Agilent Tapestation. The samples were diluted to 8 nM in Qiagen Elution Buffer (Qiagen #1014609), denatured, and loaded at ~6.4 picomolar on a Miseq at 75 base pair paired end with 8 base pair dual indexes using the MiSeq Reagent Kit v3, 150 cycles (Illumina #15043893). Resulting sequence data was used to recalculate sample concentration adjusting for sample representation. Using these updated concentrations, samples were again diluted to 8 nM in Qiagen Elution Buffer, denatured, and loaded at 6.4 picomolar on an Illumina cBOT using the HiSeq® 4000 PE Cluster Kit (Illumina #PE-410-1001). The ATAC-Seq libraries were sequenced on a HiSeq® 4000 at 75 base pair paired end with 8 base pair dual indexes using the HiSeq® 4000 SBS Kit, 150 cycles (Illumina #FC-410-1002). The sequence intensity files were generated on instrument using the Illumina Real Time Analysis software.

**ATAC-seq alignment, QC, peak calling and differential accessibility analysis**

The ATAC-seq fastq files were aligned to the hg38 reference genome downloaded from ftp://hgdownload.cse.ucsc.edu/goldenPath/hg38/bigZips/analysisSet/ on December 7, 2017 using bowtie2 v2.3.4.1 (2) and then sorted using samtools v1.8 (3). Duplicates were marked and removed using Picard MarkDuplicates v2.18.7. Low quality mapped reads (below 20) were removed using samtools, and peaks and their summits were called using macs2 v2.1.1 (4), with a p-value threshold cutoff of 0.01. We generated a bigwig file for read density visualization using DeepTools 3.1.0 (5) in 10 base pair (bp) bins using RPKM normalization. We selected 161,539 100 bp regions for a transcription factor (TF)-binding site analysis using R v3.5.0. Briefly, we used the R package DiffBind v2.8.0 (6) to identify all overlapping peaks found in at least two samples, and recentered each overlapping peak by the summit average. For each sample, we next used DiffBind to obtain the number of reads that mapped to these newly-identified regions. We identified the regions showing differential accessibility between treatment with BRM011 and control (DMSO) using edgeR v3.22.3 (7,8). We selected the genome regions showing significantly more (9362 regions) and less (13,470 regions) accessibility after treatment using a Bonferonni-corrected cutoff of 0.05. We identified TF-binding motifs enriched in the more and less accessible regions compared to HOMER-selected GC% matched background regions using HOMER findMotifsGenome v4.9.1 (9). We annotated the genomic context of all peaks using the R bioconductor package ChIPpeakAnno v3.14.0 (10), and assigned the promoter genomic context to peaks whose center was found within 1000 bp upstream of a gene. For the promoter accessibility analyses, we selected 2500 bp peaks centered at 47,299 transcription start site (TSS) regions for further analysis, used samtools v1.8 to obtain the number of reads that mapped to these TSS regions, and used edgeR v3.22.3 for the differential analyses.

**Western blot**

Protein was extracted with 1x RIPA buffer (Boston BioProducts) supplemented with PhosSToP phosphatase inhibitor tablet (Roche) and Complete protease inhibitor tablet (Roche). Quantification was performed using DC Protein Assay (Bio-Rad). 30-45 µg of protein/sample (diluted in NuPage LDS Sample buffer 4x, Thermo Fisher Scientific) was run on Criterion TGX gels (Bio-Rad) at 150V. Samples were transferred onto nitrocellulose using a TransBlot Turbo apparatus (Bio-Rad) on the mixed molecular weight setting. Membranes were blocked with 5% milk-TBS-T (Teknova) for 1 hour at room temperature followed by a 4°C overnight incubation with primary antibody diluted in 5% milk-TBS-T. Blots were washed 4x 10 minutes with TBS-T (rocking at room temperature) followed by incubation with secondary antibody goat anti-rabbit or goat anti-mouse diluted 1:10,000 in 5% milk-TBS-T for 1 hour at room temperature. Washes were repeated and membranes were then incubated SuperSignal West Femto Chemiluminescent Substrate (Thermo Fisher Scientific) for 5 min before imaging on a ChemiDoc gel imaging system (Bio-Rad).

**RT-qPCR from *in vivo* samples**

Lysis buffer from Qiagen RNeasy plus mini kit was added to tumors samples, which were then homogenized using Lysing Matrix D beads using a Precellys 24 homogenizer. Samples were then further homogenized by passing through Qiashredder columns (Qiagen), and RNA was isolated using the Qiagen RNeasy plus mini kit according to manufacturer’s protocol. cDNA synthesis was performed according to manufacturer’s protocol using ABI high capacity cDNA synthesis kit and 1 µg input RNA from each tumor. RT-qPCR was performed in technical duplicate (2 qPCR reactions per tumor sample) with FastStart Universal Probe mastermix with Rox (Roche) using a CFX384 Touch™ Real-Time PCR Detection System (Bio-Rad). Relative quantification for each sample was calculated using the 2-ΔΔCt method, normalized to β-actin, and expressed as fold change relative to vehicle control for each experiment. Graphing and analyses were performed using GraphPad Prism 7 (Graphpad Software).

**Washout experiment**

MV4-11 cells were plated and dosed with 300 nM BRM014 or DMSO for 7, 17, 24 or 48 h. For the washout samples, after 24 h treatment, cells were pelleted and washed 3 times with PBS then resuspended in media without compound and incubated for an additional 7, 17 or 24 h. Cells were then pelleted and RNA was isolated using the Qiagen RNeasy mini kit according to the manufacturer’s protocol. cDNA was synthesized using the ABI High Capacity cDNA Reverse Transcription kit according to the manufacturer’s protocol and qPCR was run as above for the *in vivo* samples.
