## Supplementary Figured and Tables for "Exquisite Sensitivity to Dual BRG1/BRM ATPase Inhibitors Reveals Broad SWI/SNF Dependencies in Acute Myeloid Leukemia"

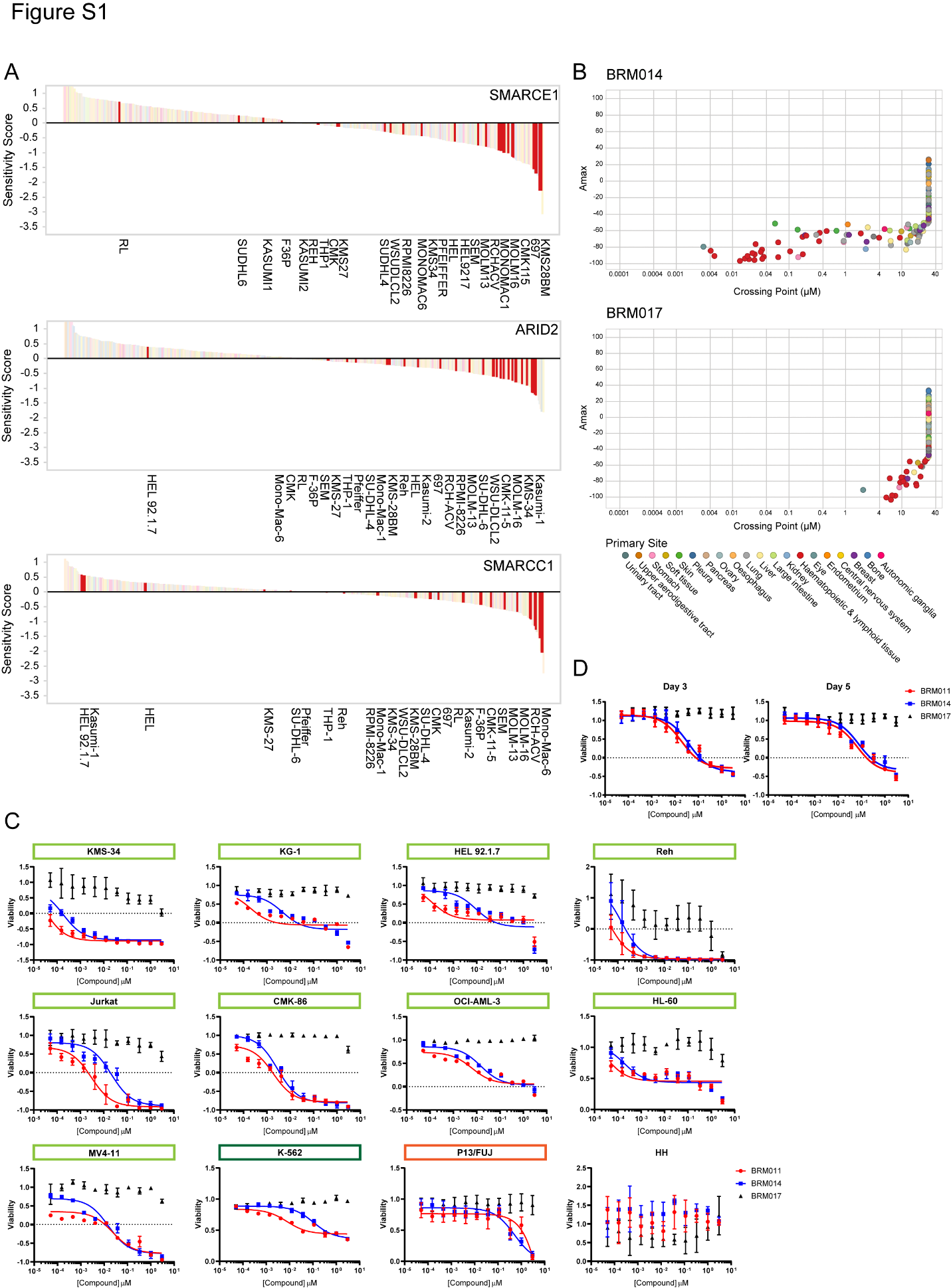

**Supplementary Figure 1: The hematopoietic lineage is dependent on SWI/SNF complex components for survival** (A) DRIVE pooled shRNA screen data (20) showing sensitivity of cancer cell lines to knockdown of SMARCE1, ARID2, and SMARCC1. Data are plotted as sensitivity score for indicated genes (y) across all cell lines (x). Hematopoietic cancer cell lines are highlighted in red and labeled. (B) Profiling of100 cancer cell lines treated with BRM014 (top) or BRM017 (bottom) in 3-day CTG proliferation assays. Amax (y) is plotted versus crossing point (x) for each cell line. Cell lines are colored by primary site as indicated. Cell line identities together with Amax and crossing point values are listed in Supplementary Table S3. (C) 12-point dose response curves for indicated hematopoietic cells treated with BRM011, BRM014, and BRM017 (N=3 per treatment, error bars shown as s.d.). Cell lines were categorized as follows: highly responsive AAC_50_ < 10 nM (light green), moderately responsive 10 nM < AAC50 < 100 nM (dark green), weakly responsive AAC50 > 100 nM (orange). (D) 12-point dose response curves for human PBMCs treated with BRM011, BRM014, and BRM017 (N=3 per treatment, error bars shown as s.d.).

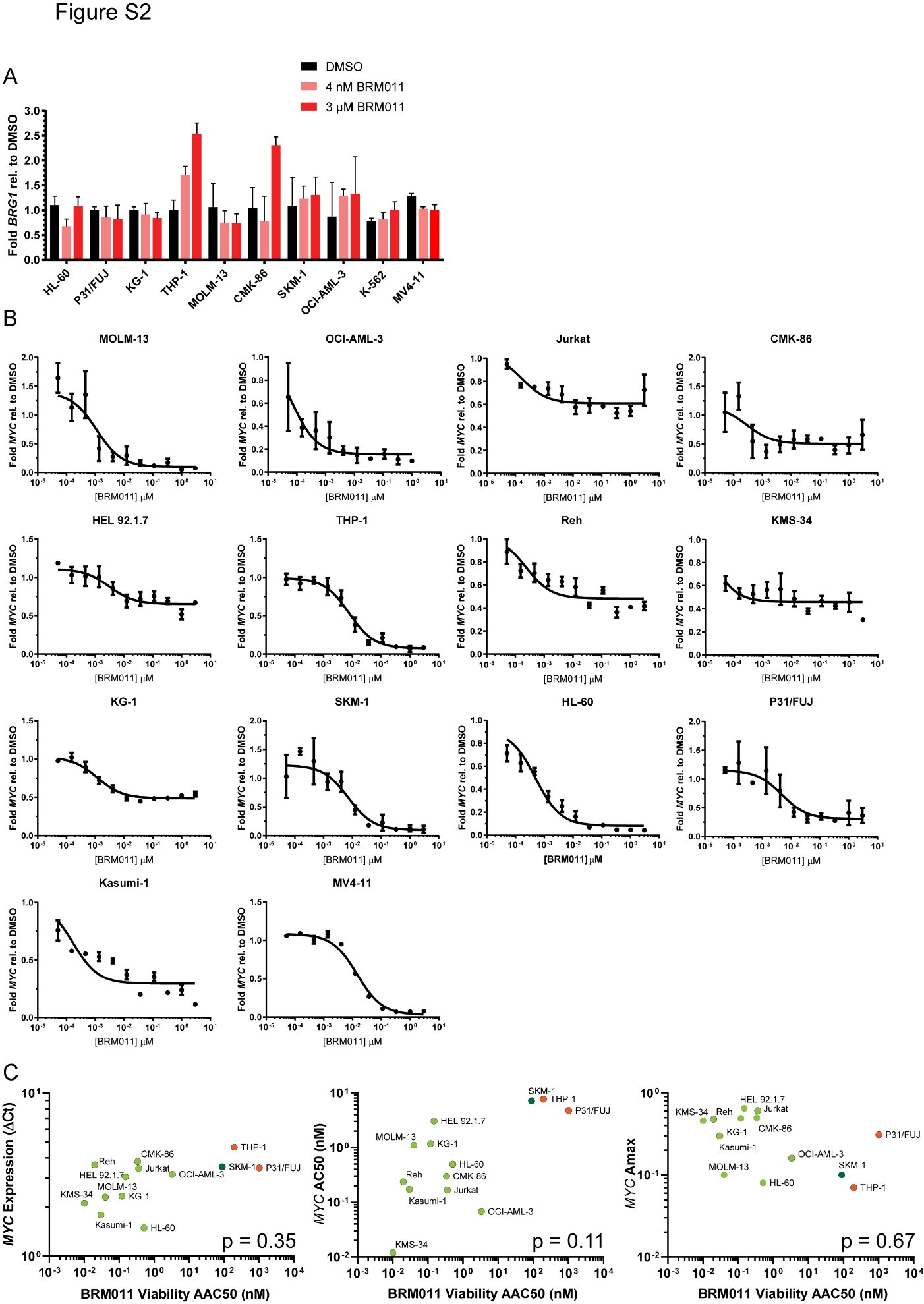

**Supplementary Figure 2: Effects of BRM011 on MYC and BRG1 gene expression.** (A) RT-qPCR analysis of *BRG1* expression in hematopoietic cancer cell lines treated with DMSO, 0.004 µM or 3 µM BRM011 for 24 hours. Ct values were normalized to *ACTB* and plotted as fold change relative to DMSO (N=3, error bars shown as s.d.). (B) 11-point MYC qPCR dose response curve for AML cell lines treated with BRM011 (N=3, error bars shown as s.d.). (C) BRM011 viability AAC50 is plotted versus basal *MYC* expression or BRM011-treated AC50 or Amax is plotted. Correlation was tested using the Spearman r test; the resulting p-value is shown for each graph.

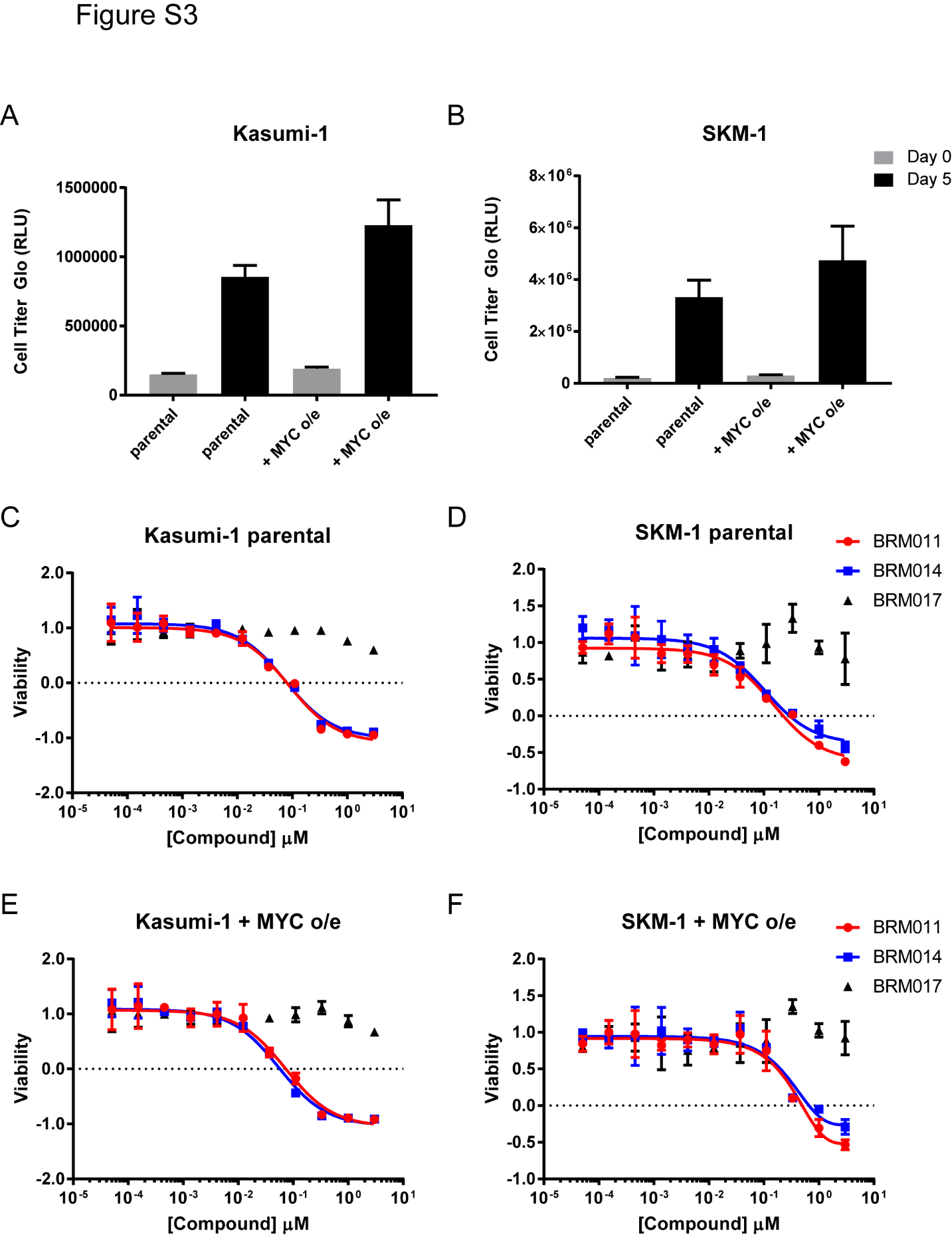

**Supplementary Figure 3: *MYC* overexpression does not affect sensitivity to SWI/SNF inhibition.** (A-B) Proliferation measured by Cell Titer Glo is plotted at 0 and 5 days in the indicated cell lines. (N=3 per condition, error bars shown as s.d.) (C-F) 5-day proliferation assay of Kasumi-1 (C) and SKM-1 (D) parental and MYC overexpression cell lines (E and F for Kasumi-1 and SKM-1, respectively) treated with BRM011, BRM014 and BRM017 (N=3 per condition, error bars shown as s.d.) BRM011 data replotted from Figure 3C-D.

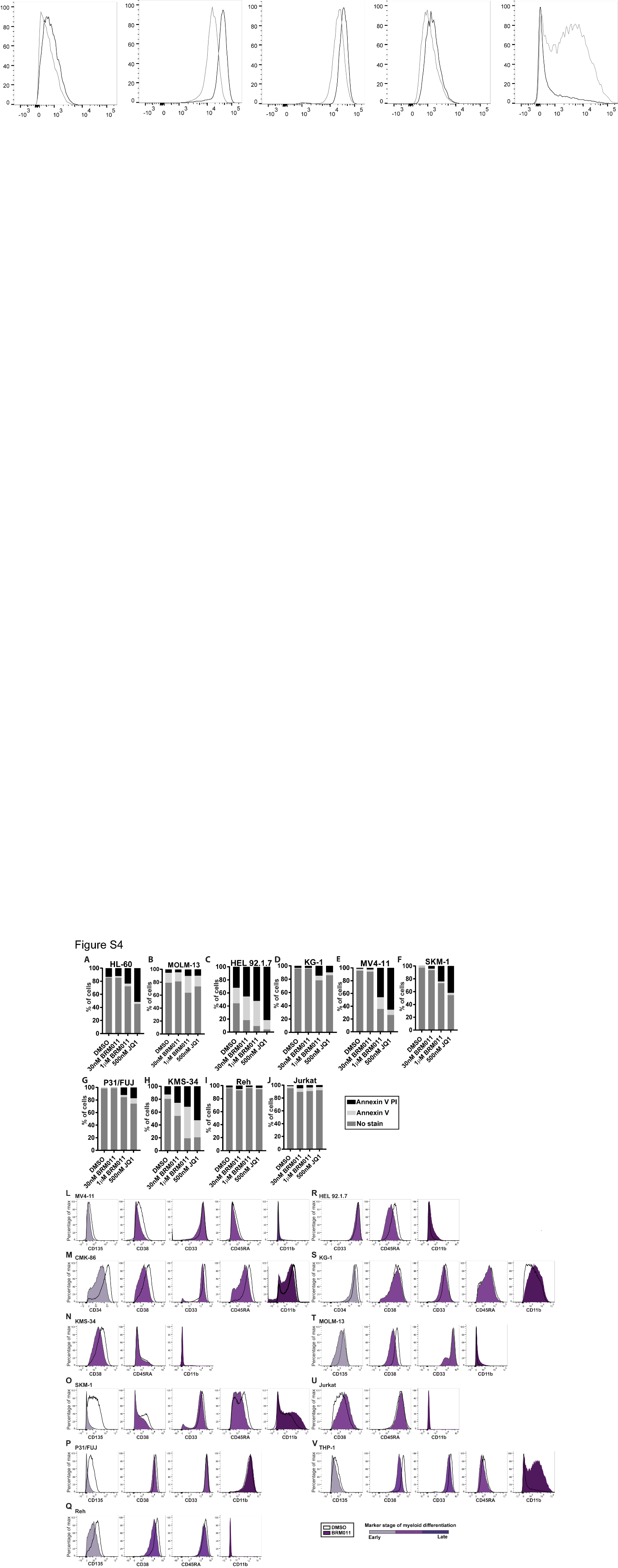

**Supplementary Figure 4: Treatment with BRM011 induces apoptosis broadly across AML cell lines but leads to differentiation in only a subset of cell lines**

(A-J) Percentage of cells unstained, Annexin V stained or Annexin V and propidium iodide (PI) stained as measured by flow cytometry after 48 hours in DMSO or labeled doses of BRM011. 500 nM JQ1 was used as a positive control. (L-V) Flow cytometry data from cells treated with DMSO (white histograms), 10 nM BRM011 (S, purple histograms), 30nM BRM011 (L-N, P-R, T, U purple histograms), 200 nM BRM011 (O, purple histograms), or 300 nM BRM011 (V, purple histograms) for 72 hours. Cells were stained with antibodies against CD34, CD135, CD38, CD33, CD45RA and CD11b.

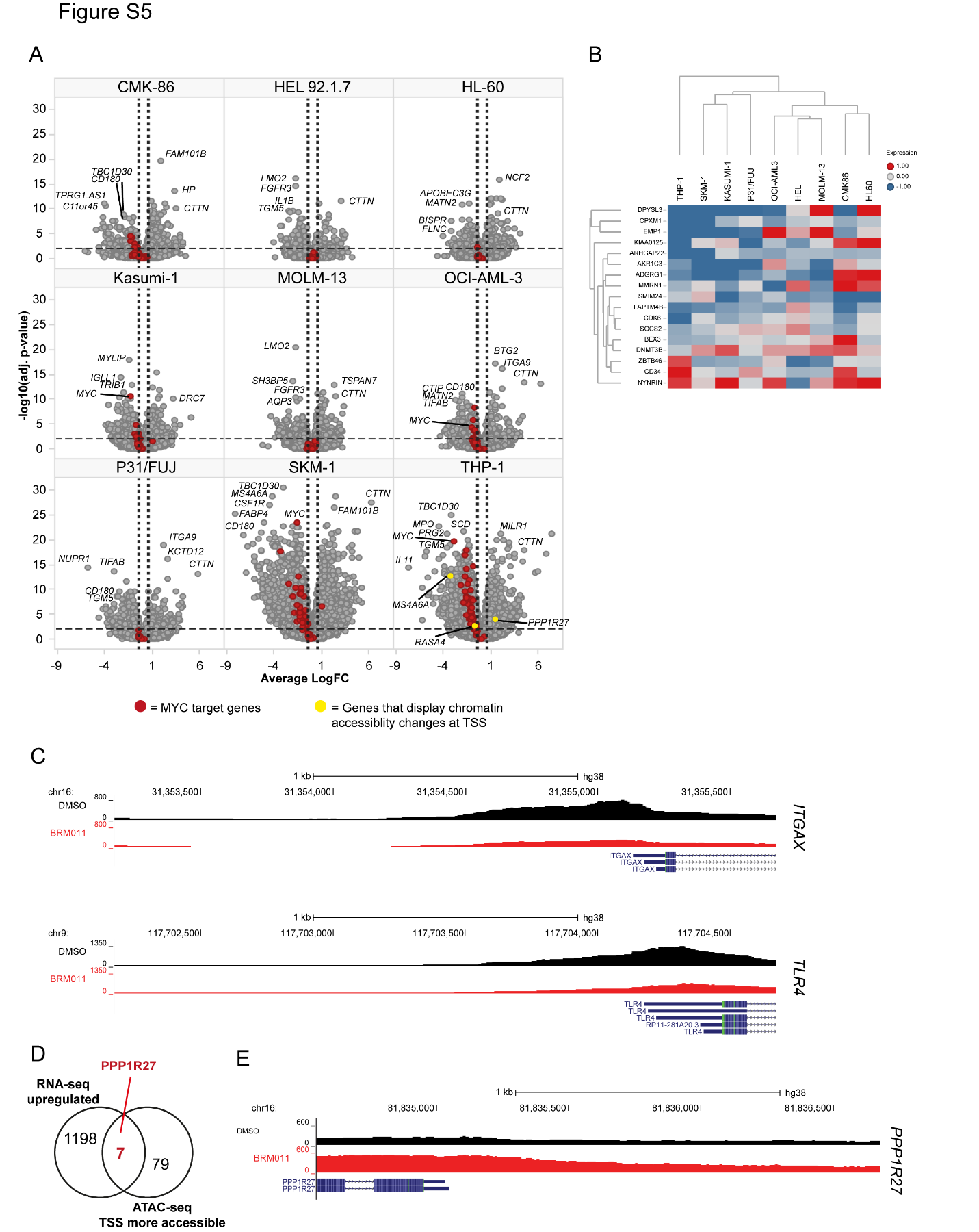

**Supplementary Figure 5: Modulation of gene expression and chromatin accessibility across AML cell lines.** (A) Gene expression changes in AML cells treated with BRM011 for 24 hours. Volcano plots depict average log fold change (logFC) in gene expression for BRM011 treatment relative to DMSO (x) versus significance (-log10[adjusted p-value]) (y); p<0.01 was considered significant (N=3 replicates/condition). (B) Heatmap of 17-gene LSC signature (23,23) depicting logFC in expression of these genes in response to BRM011 treatment across the AML cell line panel. (C) Examples of less accessible regions at gene promoters following BRM011 treatment as measured by ATAC-Seq. (D) Comparison of RNA-seq downregulated genes and ATAC-seq TSS less accessible genes. (E) Example of more accessible regions at gene promoters following BRM011 treatment as measured by ATAC-Seq.

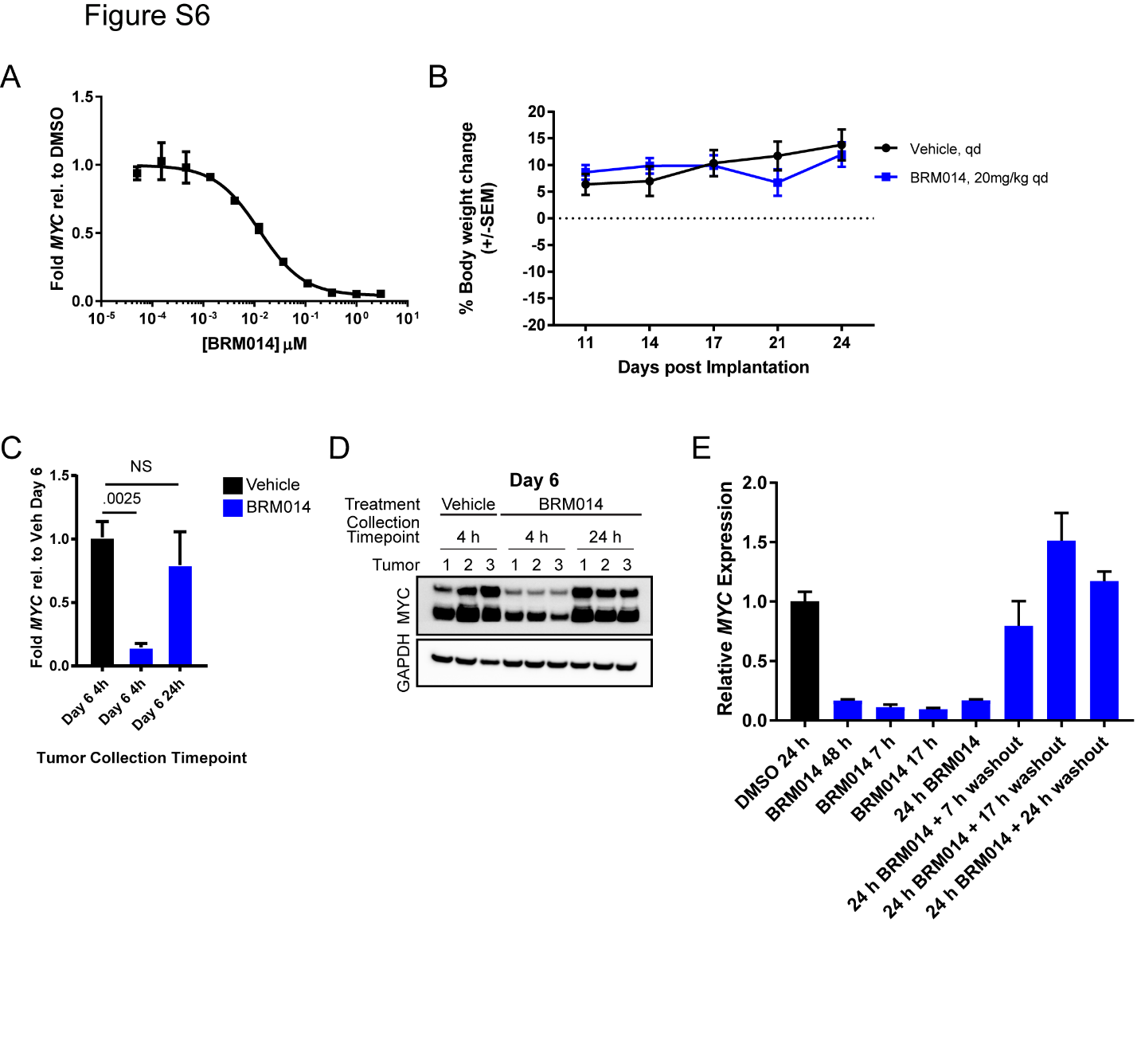

**Supplementary Figure 6: MYC repression is not maintained throughout SWI/SNF inhibitor dosing *in vivo.*** (A) RT-qPCR analysis of *MYC* expression in MV4-11 cells treated with BRM014 for 24 hours. Expression was normalized to *ACTB* and plotted as fold change relative to DMSO (N=3, error bars shown as s.d.).(B) Body weight change for mice treated with either vehicle or BRM014 20mg/kg for 2 weeks. (C) Fold change in *MYC* expression across tumors treated with vehicle or BRM014 measured by RT-qPCR at 7-day timepoint. Plotted as average +/- SEM. N=3 per time point. Student’s two tailed t-test, p-value indicated. (D) Immunoblots of MYC and PARP levels in mice treated with vehicle or BRM014 for labeled periods of time. Each column represents tumor from a unique mouse. GAPDH was used as a loading control. (E) RT-qPCR analysis of *MYC* expression in MV-4-11 cells treated with DMSO or 300 nM BRM014 for indicated times with or without washout of BRM014 from cell culture media following treatment with compound (N=3 per condition, error bars shown as s.d.).

Supplementary Tables:

**Table S1:**

| **Cell line** | **Cell culture media** | **5-day Proliferation assay seeding density** |
| --- | --- | --- |
| CMK-86 | RPMI-1640 + 10% FBS | 3000 |
| HEL 92.1.7 | RPMI-1640 + 10% FBS | 1500 |
| HH | RPMI-1640 + 10% FBS | 3000 |
| HL-60 | IMDM + 10% FBS | 2000 |
| Jurkat | RPMI-1640 + 10% FBS | 1000 |
| K-562 | IMDM + 10% FBS | 1500 |
| Kasumi-1 | RPMI-1640 + 20% FBS | 1500 |
| KG-1 | IMDM + 10% FBS | 1000 |
| KMS-34 | RPMI-1640 + 10% FBS | 500 |
| MOLM-13 | RPMI-1640 + 10% FBS | 3000 |
| MV-4-11 | IMDM + 10% FBS | 1500 |
| OCI-AML3 | RPMI-1640 + 10% FBS | 3000 |
| P31/FUJ | RPMI-1640 + 10% FBS | 3000 |
| Reh | RPMI-1640 + 10% FBS | 4500 |
| SKM-1 | RPMI-1640 + 10% FBS | 3000 |
| THP-1 | RPMI-1640 + 10% FBS | 3000 |

**Table S1: Proliferation assay seeding conditions.**

**Table S2:**

| **Gene** | **Catalog #** | **Vendor** |
| --- | --- | --- |
| *BRG1/SMARCA4* | HS00231324 | ABI |
| *BRM/SMARCA2* | HS01030858 | ABI |
| *MYC* | HS80153408 | ABI |
| *B2M* | 4325797 | ABI |
| *β-actin* | Custom synthesis:  β-actinF: GCGAGAAGATGACCCAGATC  β-actinR: CCAGTGGTACGGCCAGAGG  β-actinHEX: VIC-CCAGCCATGTACGTTGCTATCCAGGC-HEX | IDT |

**Table S2: RT-qPCR probes used in this study.**

**Table S3:**

| Cell Line | Primary site | BRM014 Amax | BRM014 Crossing Point (µM) | BRM011 Amax | BRM011 Crossing Point (µM) | BRM017 Amax | BRM017 Crossing Point (µM) |
| --- | --- | --- | --- | --- | --- | --- | --- |
| 92.1 | eye | -79.823 | 0.003 | -88.693 | 0.001 | -91.139 | 2.073 |
| 697 | haematopoietic & lymphoid | -92.340 | 0.065 | -92.539 | 0.209 | -98.044 | 6.779 |
| 769-P | kidney | -82.387 | 2.305 | -77.813 | 2.893 | 16.869 | 30.000 |
| A172 | central nervous system | -47.138 | 30.000 | 0.522 | 30.000 | 19.553 | 30.000 |
| A2780 | ovary | -74.491 | 14.289 | -45.361 | 30.000 | -43.484 | 30.000 |
| A4/Fuk | haematopoietic & lymphoid | -87.234 | 0.023 | -90.142 | 0.038 | -74.314 | 21.487 |
| A549 | lung | -68.104 | 20.262 | -48.322 | 30.000 | -46.221 | 30.000 |
| ABC-1 | lung | -65.296 | 1.051 | -60.148 | 0.230 | -9.569 | 30.000 |
| AsPC-1 | pancreas | -49.468 | 30.000 | -44.251 | 30.000 | -32.727 | 30.000 |
| AU565 | breast | -73.767 | 2.378 | -69.962 | 3.218 | -43.780 | 30.000 |
| BT-549 | breast | -54.069 | 24.599 | -21.124 | 30.000 | 4.678 | 30.000 |
| C3A | liver | -60.687 | 23.936 | -21.217 | 30.000 | -48.420 | 30.000 |
| Caki-2 | kidney | 13.174 | 30.000 | -1.035 | 30.000 | 12.697 | 30.000 |
| CAL 27 | upper aerodigestive tract | -47.067 | 30.000 | -55.472 | 18.354 | 4.628 | 30.000 |
| CAL-120 | breast | -32.642 | 30.000 | -46.982 | 30.000 | 16.308 | 30.000 |
| CAL-29 | urinary tract | -28.971 | 30.000 | -28.839 | 30.000 | 12.227 | 30.000 |
| CCRF-HSB-2 | haematopoietic & lymphoid | -60.746 | 0.466 | -52.222 | 1.852 | -51.825 | 28.300 |
| CMK-86 | haematopoietic & lymphoid | -91.033 | 0.018 | -89.944 | 0.029 | -76.633 | 6.147 |
| COLO-704 | ovary | -55.943 | 21.676 | -41.300 | 30.000 | -11.894 | 30.000 |
| DB | haematopoietic & lymphoid | -66.385 | 0.204 | -62.737 | 0.479 | -56.000 | 25.855 |
| DMS 53 | lung | -72.511 | 0.939 | -87.374 | 0.089 | -29.174 | 30.000 |
| DND-41 | haematopoietic & lymphoid | -92.055 | 0.036 | -92.393 | 0.030 | -54.840 | 26.055 |
| EOL-1 | haematopoietic & lymphoid | -88.962 | 0.025 | -87.890 | 0.010 | -81.620 | 9.383 |
| FU97 | stomach | -92.344 | 0.147 | -95.041 | 0.158 | -87.766 | 9.082 |
| G-402 | soft tissue | -61.126 | 9.919 | -31.554 | 30.000 | -56.921 | 19.471 |
| GDM-1 | haematopoietic & lymphoid | -94.764 | 0.020 | -95.531 | 0.010 | -100.529 | 5.320 |
| GP2d | large intestine | 9.053 | 30.000 | 12.244 | 30.000 | 11.695 | 30.000 |
| GRANTA-519 | haematopoietic & lymphoid | -62.733 | 0.352 | -57.509 | 0.280 | -37.459 | 30.000 |
| HCC1500 | breast | -42.893 | 30.000 | -48.501 | 30.000 | -5.397 | 30.000 |
| HCC1806 | breast | -59.577 | 21.033 | -29.935 | 30.000 | -47.527 | 30.000 |
| HCC1954 | breast | -40.428 | 30.000 | -40.614 | 30.000 | -30.139 | 30.000 |
| HCC-44 | lung | -15.044 | 30.000 | -34.529 | 30.000 | 9.318 | 30.000 |
| HCC827 | lung | -36.726 | 30.000 | -30.048 | 30.000 | 9.049 | 30.000 |
| HEC-108 | endometrium | -43.739 | 30.000 | -2.872 | 30.000 | 5.443 | 30.000 |
| HEC-251 | endometrium | -64.675 | 18.563 | -17.137 | 30.000 | 12.957 | 30.000 |
| Hep 3B2.1-7 | liver | -72.511 | 19.316 | -40.473 | 30.000 | -2.108 | 30.000 |
| HPB-ALL | haematopoietic & lymphoid | -76.388 | 0.026 | -74.595 | 0.047 | -55.542 | 13.843 |
| Hs 936.T | skin | -51.634 | 0.057 | -56.350 | 0.075 | -19.773 | 30.000 |
| HS-SY-II |  | -41.224 | 30.000 | -58.684 | 0.646 | -40.182 | 30.000 |
| HT-1080 | soft tissue | 4.876 | 30.000 | -67.860 | 0.438 | -15.061 | 30.000 |
| HT-29 | large intestine | -50.763 | 24.654 | -26.998 | 30.000 | -27.701 | 30.000 |
| HuH-6 | liver | -72.645 | 6.342 | -55.293 | 9.863 | -68.873 | 16.683 |
| HuH-7 | liver | -83.281 | 11.853 | -19.901 | 30.000 | -49.733 | 30.000 |
| HUP-T4 | pancreas | -47.554 | 30.000 | -52.375 | 0.860 | -21.234 | 30.000 |
| IPC-298 | skin | -60.439 | 0.696 | -65.493 | 3.252 | -25.201 | 30.000 |
| JHOM-1 | ovary | -38.430 | 30.000 | 4.534 | 30.000 | 25.561 | 30.000 |
| JHOS-2 | ovary | -27.790 | 30.000 | -32.979 | 30.000 | 0.663 | 30.000 |
| JL-1 | pleura | 20.586 | 30.000 | 9.183 | 30.000 | 3.216 | 30.000 |
| KBM-7 | haematopoietic & lymphoid | -63.055 | 17.609 | -28.808 | 30.000 | -19.832 | 30.000 |
| KE-97 | haematopoietic & lymphoid | -83.173 | 0.033 | -83.623 | 0.039 | -47.354 | 30.000 |
| KNS-42 | central nervous system | -22.355 | 30.000 | -19.432 | 30.000 | 14.722 | 30.000 |
| KO52 | haematopoietic & lymphoid | -67.720 | 0.174 | -64.924 | 0.265 | 2.197 | 30.000 |
| KP4 | pancreas | -38.507 | 30.000 | -24.510 | 30.000 | 15.854 | 30.000 |
| KYSE-180 | oesophagus | -46.015 | 30.000 | -1.180 | 30.000 | -34.339 | 30.000 |
| LP-1 | haematopoietic & lymphoid | -73.033 | 0.343 | -66.779 | 1.157 | -83.529 | 17.988 |
| LS 180 | large intestine | -50.450 | 27.232 | -52.660 | 3.119 | -1.755 | 30.000 |
| MDA-MB-453 | breast | -65.465 | 0.195 | -67.786 | 0.085 | -76.881 | 12.641 |
| MEL-HO | skin | -58.874 | 0.142 | -58.433 | 0.166 | 21.748 | 30.000 |
| MFE-319 | endometrium | -52.677 | 1.104 | -59.144 | 2.249 | 7.428 | 30.000 |
| Mino | haematopoietic & lymphoid | -96.867 | 0.014 | -96.391 | 0.006 | -98.363 | 7.703 |
| MOLM-13 | haematopoietic & lymphoid | -90.950 | 0.005 | -92.889 | 0.004 | -80.701 | 10.759 |
| MOLP-8 | haematopoietic & lymphoid | -74.370 | 0.065 | -68.785 | 0.037 | -68.253 | 12.212 |
| MV-4-11 | haematopoietic & lymphoid | -94.202 | 0.017 | -96.855 | 0.011 | -84.790 | 12.028 |
| NCI-H1299 | lung | -42.057 | 30.000 | -53.077 | 6.580 | -30.073 | 30.000 |
| NCI-H146 | lung | -65.028 | 21.051 | -19.306 | 30.000 | 33.685 | 30.000 |
| NCI-H1568 | lung | 8.013 | 30.000 | -11.293 | 30.000 | 1.730 | 30.000 |
| NCI-H1581 | lung | -73.639 | 19.279 | -2.053 | 30.000 | -62.137 | 23.142 |
| NCI-H1650 | lung | -34.868 | 30.000 | -1.557 | 30.000 | -16.694 | 30.000 |
| NCI-H1693 | lung | -48.406 | 30.000 | -54.163 | 0.021 | -12.944 | 30.000 |
| NCI-H1792 | lung | -47.823 | 30.000 | 2.854 | 30.000 | -34.227 | 30.000 |
| NCI-H2110 | lung | -62.280 | 16.527 | 6.045 | 30.000 | -41.770 | 30.000 |
| NCI-H2122 | lung | -77.841 | 6.410 | -87.315 | 4.710 | -61.576 | 22.223 |
| NCI-H2171 | lung | -1.635 | 30.000 | -11.425 | 30.000 | -1.498 | 30.000 |
| NCI-H2172 | lung | -57.843 | 23.238 | -19.561 | 30.000 | -41.915 | 30.000 |
| NCI-H2347 | lung | -52.725 | 12.991 | -72.579 | 1.834 | -40.130 | 30.000 |
| NCI-H3255 | lung | -49.854 | 30.000 | -42.567 | 30.000 | -31.770 | 30.000 |
| NCI-H441 | lung | -37.575 | 30.000 | -24.705 | 30.000 | -15.343 | 30.000 |
| NCI-H460 | lung | -51.753 | 28.529 | -35.840 | 30.000 | -35.000 | 30.000 |
| NCI-H647 | lung | -23.317 | 30.000 | 9.758 | 30.000 | -30.171 | 30.000 |
| NCI-H650 | lung | 1.008 | 30.000 | -0.033 | 30.000 | -3.890 | 30.000 |
| NCI-H810 | lung | -8.412 | 30.000 | -32.430 | 30.000 | 3.918 | 30.000 |
| NCI-H838 | lung | -45.900 | 30.000 | 7.769 | 30.000 | -18.390 | 30.000 |
| NCI-N87 | stomach | -74.167 | 0.941 | -72.963 | 3.564 | -1.233 | 30.000 |
| OCI-AML3 | haematopoietic & lymphoid | -85.884 | 0.032 | -88.242 | 0.035 | -103.483 | 6.469 |
| OUMS-23 | large_intestine | -39.003 | 30.000 | 4.146 | 30.000 | 23.408 | 30.000 |
| P12-ICHIKAWA | haematopoietic & lymphoid | -88.759 | 0.177 | -82.249 | 0.338 | -61.333 | 21.912 |
| P31/FUJ | haematopoietic & lymphoid | -63.558 | 1.878 | -55.575 | 5.254 | 12.529 | 30.000 |
| Panc 10.05 | pancreas | -49.382 | 30.000 | -54.236 | 1.620 | 9.876 | 30.000 |
| PE/CA-PJ34 (clone C12) | upper aerodigestive tract | 25.226 | 30.000 | 1.099 | 30.000 | 2.968 | 30.000 |
| RD | soft tissue | 10.837 | 30.000 | -3.478 | 30.000 | 9.359 | 30.000 |
| RERF-LC-AI | lung | -65.316 | 0.303 | -57.324 | 0.224 | -9.494 | 30.000 |
| RERF-LC-MS | lung | 26.144 | 30.000 | -0.303 | 30.000 | 12.104 | 30.000 |
| RKO | large intestine | -59.939 | 6.119 | -54.589 | 5.369 | 22.213 | 30.000 |
| RMUG-S | ovary | 14.855 | 30.000 | -4.283 | 30.000 | 10.379 | 30.000 |
| RVH-421 | skin | 4.878 | 30.000 | -4.360 | 30.000 | -7.637 | 30.000 |
| SEM | haematopoietic & lymphoid | -68.908 | 3.974 | -65.875 | 3.270 | -18.286 | 30.000 |
| SK-ES-1 | bone | -10.756 | 30.000 | 9.051 | 30.000 | 32.897 | 30.000 |
| SK-HEP-1 | liver | -24.110 | 30.000 | -47.736 | 30.000 | -2.690 | 30.000 |
| SKM-1 | haematopoietic & lymphoid | -85.194 | 0.105 | -88.505 | 0.061 | -47.564 | 30.000 |
| SK-MEL-2 | skin | 5.387 | 30.000 | 4.787 | 30.000 | 9.408 | 30.000 |
| SK-MEL-2 |  | 5.387 | 30.000 | 4.787 | 30.000 | 9.408 | 30.000 |
| SK-N-SH | autonomic ganglia | -58.422 | 4.984 | -18.261 | 30.000 | 4.538 | 30.000 |
| SNG-M | endometrium | 14.429 | 30.000 | -6.933 | 30.000 | -0.297 | 30.000 |
| SNU-16 | stomach | -56.059 | 12.364 | -54.213 | 14.125 | 10.925 | 30.000 |
| SNU-423 | liver | 6.458 | 30.000 | 0.894 | 30.000 | 0.838 | 30.000 |
| SNU-878 | liver | -56.037 | 2.490 | -58.348 | 22.899 | -31.249 | 30.000 |
| SNU-886 | liver | -20.642 | 30.000 | -16.067 | 30.000 | -3.490 | 30.000 |
| SU.86.86 | pancreas | -22.067 | 30.000 | 4.409 | 30.000 | -20.075 | 30.000 |
| SU-DHL-4 | haematopoietic & lymphoid | -94.284 | 0.034 | -93.835 | 0.025 | -76.044 | 10.670 |
| SU-DHL-5 | haematopoietic & lymphoid | -95.943 | 0.025 | -97.595 | 0.010 | -85.636 | 18.310 |
| SU-DHL-8 | haematopoietic & lymphoid | -84.686 | 0.004 | -84.381 | 0.003 | -75.479 | 9.880 |
| SUM185PE | breast | -46.090 | 30.000 | -38.325 | 30.000 | -46.574 | 30.000 |
| SUM52PE | breast | -65.297 | 1.160 | -67.964 | 1.241 | -27.865 | 30.000 |
| SW 1088 | central nervous system | -35.225 | 30.000 | -0.971 | 30.000 | 5.409 | 30.000 |
| SW 1271 | lung | -26.253 | 30.000 | -7.324 | 30.000 | -26.903 | 30.000 |
| SW480 | large intestine | -54.788 | 17.561 | -43.464 | 30.000 | 12.541 | 30.000 |
| SW837 | large intestine | -71.942 | 12.535 | -58.976 | 3.670 | -28.947 | 30.000 |
| SYO-1 | soft tissue | -66.218 | 14.419 | 9.363 | 30.000 | -29.908 | 30.000 |
| TE-14 | oesophagus | -3.102 | 30.000 | -44.872 | 30.000 | 32.112 | 30.000 |
| TE-15 | oesophagus | -47.037 | 30.000 | -41.373 | 30.000 | -39.020 | 30.000 |
| THP-1 | haematopoietic & lymphoid | -68.938 | 9.326 | -69.405 | 4.017 | -43.595 | 30.000 |
| UACC-257 | skin | -57.438 | 13.661 | -42.508 | 30.000 | -39.962 | 30.000 |
| UCH-1 | bone | -41.484 | 30.000 | 13.557 | 30.000 | 27.421 | 30.000 |
| WM1799 | skin | -18.290 | 30.000 | -21.526 | 30.000 | 16.678 | 30.000 |

**Table S3: Calculated Amax and crossing point values corresponding to curves in Figure 1B.**

**Table S4:**

| **Cell line** | **Mutations^a^** | **Subclassification** |
| --- | --- | --- |
| KMS-34 |  | Plasma cell myeloma |
| Reh | ETV6-RUNX1 gene fusion | ALL |
| Kasumi-1 | RUNX1-RUNX1T1 gene fusion  Het. KIT N822K  RAD21 K330Profs*6  Hom. TP53 R248Q | AML |
| MOLM-13 | MLL-AF9 gene fusion  FLT3-ITD | AML |
| KG-1 | FGFR1OP2-FGFR1 gene fusion  NRAS G12D  Hom. TP53 p.672+1 G>A | AML |
| HEL 92.1.7 | Hom. JAK2 V617F  Hom. TP53 M133K | AML |
| CMK-86 | TP53-FXR2 gene fusion  Het. TP53 D49H | AML |
| Jurkat | Het. BAX E41Rfs and E41Gfs  Het. FBZW7 R505C  Het. INPP5D Q345*  Hom. MSH2 R711*  Hom. MSH6 F1088Sfs*2  Het. NOTCH1 R1627H  Het. TP53 R196* | T cell leukemia |
| HL-60 | Hom. CDKN2A R80*  Het. NRAS Q61L  Hom. TP53 del | AML |
| OCI-AML-3 | DNMT3A R882C  Het. NPM1 W288Cfs*12 | AML |
| K-562 | BCR-ABL1 gene fusion  Hom. TP53 Q136fs*13 | CML |
| SKM-1 | Hom. KRAS K117N  Hom. TP53 R248Q | AML |
| THP-1 | CSNK2A1-DDX39B gene fusion  MLL-AF9 gene fusion  Het. NRAS G12D  Het TP53 R174fs*3 | AML |
| P31/FUJ | Het. KMT2A R158fs*13  Het. NRAS G12D  Het. PTEN E7*  Het. RAD21 H208R  Het. TP53 R196* | AML |
| HH | FOXK2-TP63 gene fusion  Hom. TP53 c.376-1G>A splice acceptor mut. | Cutaneous T cell lymphoma |
| MV4-11 | MLL-AFF1 gene fusion  FLT3-ITD |  |

^a^ mutation information taken from Cellosaurus annotations (https://web.expasy.org/cellosaurus/)

**Table S4: Mutation annotation for cell line panel.**

**Table S5:**

| **Cell line** | **Compound** | **AAC50 (nM)** | **Amax** | **R^2^** |
| --- | --- | --- | --- | --- |
| KMS-34 | BRM011 | 0.01 | -0.89 | 0.97 |
|  | BRM014 | 0.04 | -0.85 | 0.94 |
| Reh | BRM011 | 0.02 | -0.96 | 0.94 |
|  | BRM014 | 0.09 | -1.00 | 0.89 |
| Kasumi-1 | BRM011 | 0.03 | -0.93 | 0.96 |
|  | BRM014 | 0.3 | -0.96 | 0.97 |
| MOLM-13 | BRM011 | 0.04 | -0.10 | 0.78 |
|  | BRM014 | 0.5 | -0.09 | 0.85 |
| KG-1 | BRM011 | 0.1 | -0.06 | 0.90 |
|  | BRM014 | 1.7 | -0.18 | 0.93 |
| HEL 92.1.7 | BRM011 | 0.2 | 0.07 | 0.89 |
|  | BRM014 | 5.4 | -0.12 | 0.92 |
| CMK-86 | BRM011 | 0.3 | -0.79 | 0.93 |
|  | BRM014 | 1.1 | -0.81 | 0.97 |
| Jurkat | BRM011 | 0.4 | -0.92 | 0.93 |
|  | BRM014 | 4.6 | -0.89 | 0.95 |
| HL-60 | BRM011 | 0.5 | 0.46 | 0.88 |
|  | BRM014 | 1.5 | 0.43 | 0.89 |
| MV4-11 | BRM011 | ND | -0.77 | 0.80 |
|  | BRM014 | 2.4 | -0.78 | 0.94 |
| OCI-AML-3 | BRM011 | 3.3 | 0.06 | 0.87 |
|  | BRM014 | 11.3 | 0.03 | 0.96 |
| K-562 | BRM011 | 42.0 | 0.44 | 0.90 |
|  | BRM014 | 296.9 | 0.36 | 0.98 |
| SKM-1 | BRM011 | 88.5 | -0.65 | 0.81 |
|  | BRM014 | 50.2 | -0.72 | 0.97 |
| THP-1 | BRM011 | 195.6 | 0.40 | 0.77 |
|  | BRM014 | 332.0 | 0.23 | 0.96 |
| P31/FUJ | BRM011 | 1011.6 | ND | 0.70 |
|  | BRM014 | 358.2 | -0.01 | 0.88 |
| HH | BRM011 | ND | ND | - |
|  | BRM014 | ND | ND | - |
| PBMC^a^ | BRM011 | 35 | -0.40 | 0.93 |
|  | BRM014 | 51 | -0.33 | 0.95 |

^a^ data fit from Day 5 viability curves

**Table S5: Calculated proliferation inhibition values corresponding to curves in Figure 1C, S1C, S1D.**

**Table S6:**

| **Cell line** | **Compound** | **AC50 (nM)** | **Amax** | **R^2^** |
| --- | --- | --- | --- | --- |
| KMS-34 | BRM011 | 0.01 | 0.46 | 0.93 |
| Reh | BRM011 | 0.2 | 0.48 | 0.85 |
| Kasumi-1 | BRM011 | 0.2 | 0.30 | 0.83 |
| MOLM-13 | BRM011 | 1.0 | 0.10 | 0.80 |
| KG-1 | BRM011 | 1.0 | 0.49 | 0.96 |
| HEL 92.1.7 | BRM011 | 3.0 | 0.65 | 0.91 |
| CMK-86 | BRM011 | 0.3 | 0.50 | 0.49 |
| Jurkat | BRM011 | 0.2 | 0.61 | 0.85 |
| HL-60 | BRM011 | 0.5 | 0.08 | 0.86 |
| MV4-11 | BRM011 | 14.6 | 0.03 | 0.98 |
| OCI-AML-3 | BRM011 | 0.07 | 0.16 | 0.74 |
| SKM-1 | BRM011 | 7.0 | 0.10 | 0.81 |
| THP-1 | BRM011 | 8.0 | 0.07 | 0.97 |
| P31/FUJ | BRM011 | 5.0 | 0.31 | 0.77 |

**Table S6: Calculated *MYC* inhibition values.**

**Table S7:**

|  |  | **Basal Markers Expressed** | | |  |  |
| --- | --- | --- | --- | --- | --- | --- |
| **Cell line** | **CD34** | **CD135** | **CD38** | **CD33** | **CD45RA** | **CD11b** |
| **Kasumi** | Y | N | Y | Y | Y | N |
| **MV411** | N | Y | Y | Y | Y | N |
| **KMS34** | N | N | Y | N | Y | N |
| **KG-1** | Y | N | Y | Y | Y | Y |
| **HEL** | N | N | N | Y | Y | N |
| **Molm13** | N | Y | Y | Y | N | N |
| **HL60** | N | Y | Y | Y | Y | N |
| **CMK86** | Y | N | Y | Y | Y | Y |
| **OCI-AML3** | N | Y | Y | Y | Y | N |
| **SKM1** | N | Y | Y | Y | Y | Y |
| **P31/FUJ** | N | Y | Y | Y | N | Y |
| **THP1** | N | Y | Y | Y | Y | Y |
| **REH** | N | Y | Y | N | Y | N |
| **Jurkat** | N | N | Y | N | Y | N |

**Table S7: Expression of cell surface markers across cell line panel.**

**Table S8:**

| Pathway | CMK-86 | HEL 92.1.7 | HL-60 | Kasumi-1 | MOLM-13 | OCI-AML-3 | P31/FUJ | SKM-1 | THP-1 |
| --- | --- | --- | --- | --- | --- | --- | --- | --- | --- |
| HALLMARK_ADIPOGENESIS |  |  |  |  | X | X |  | X | X |
| HALLMARK_ALLOGRAFT_REJECTION |  | X |  | X |  |  | X | X | X |
| HALLMARK_ANDROGEN_RESPONSE |  |  |  |  |  | X |  | X | X |
| HALLMARK_ANGIOGENESIS | X |  |  | X |  |  |  | X |  |
| HALLMARK_APICAL_JUNCTION | X |  |  | X |  |  |  | X |  |
| HALLMARK_APICAL_SURFACE |  |  |  |  |  |  |  |  |  |
| HALLMARK_APOPTOSIS | X |  |  | X | X | X |  | X | X |
| HALLMARK_BILE_ACID_METABOLISM |  |  |  |  |  |  |  |  | X |
| HALLMARK_CHOLESTEROL_HOMEOSTASIS |  |  |  | X |  |  |  | X | X |
| HALLMARK_COAGULATION |  |  |  |  | X |  |  | X |  |
| HALLMARK_COMPLEMENT |  |  |  |  | X |  |  | X |  |
| HALLMARK_DNA_REPAIR |  |  |  |  |  |  |  | X | X |
| HALLMARK_E2F_TARGETS |  |  |  |  |  | X |  | X | X |
| HALLMARK_EPITHELIAL_MESENCHYMAL_TRANSITION | X |  | X | X |  | X |  | X | X |
| HALLMARK_ESTROGEN_RESPONSE_EARLY | X |  | X | X |  | X |  | X | X |
| HALLMARK_ESTROGEN_RESPONSE_LATE | X |  | X | X |  | X |  | X | X |
| HALLMARK_FATTY_ACID_METABOLISM |  | X |  |  |  |  |  | X | X |
| HALLMARK_G2M_CHECKPOINT |  |  |  |  |  | X |  | X | X |
| HALLMARK_GLYCOLYSIS |  |  |  |  |  | X |  | X | X |
| HALLMARK_HEDGEHOG_SIGNALING |  |  |  |  |  |  |  | X |  |
| HALLMARK_HEME_METABOLISM | X | X |  |  | X | X |  | X | X |
| HALLMARK_HYPOXIA | X |  |  |  | X | X |  | X | X |
| HALLMARK_IL2_STAT5_SIGNALING | X |  |  | X |  | X | X | X | X |
| HALLMARK_IL6_JAK_STAT3_SIGNALING |  | X |  | X |  | X |  | X | X |
| HALLMARK_INFLAMMATORY_RESPONSE | X |  |  | X | X | X | X | X | X |
| HALLMARK_INTERFERON_ALPHA_RESPONSE |  | X |  |  |  | X | X | X | X |
| HALLMARK_INTERFERON_GAMMA_RESPONSE |  | X |  |  |  | X | X | X | X |
| HALLMARK_KRAS_SIGNALING_DN |  |  |  |  |  |  |  |  |  |
| HALLMARK_KRAS_SIGNALING_UP | X |  | X | X |  | X | X | X | X |
| HALLMARK_MITOTIC_SPINDLE |  |  |  |  | X |  |  | X | X |
| HALLMARK_MTORC1_SIGNALING | X |  |  | X | X | X |  | X | X |
| HALLMARK_MYC_TARGETS_V1 |  |  |  |  |  | X |  | X | X |
| HALLMARK_MYC_TARGETS_V2 | X |  |  |  |  | X |  | X | X |
| HALLMARK_MYOGENESIS |  |  |  | X |  |  |  | X | X |
| HALLMARK_NOTCH_SIGNALING |  |  |  | X |  |  |  | X |  |
| HALLMARK_OXIDATIVE_PHOSPHORYLATION |  |  |  |  |  | X |  | X | X |
| HALLMARK_P53_PATHWAY | X |  |  |  | X | X |  | X | X |
| HALLMARK_PANCREAS_BETA_CELLS |  |  |  |  |  |  |  |  |  |
| HALLMARK_PEROXISOME |  |  |  |  |  |  |  | X | X |
| HALLMARK_PI3K_AKT_MTOR_SIGNALING |  |  |  |  |  |  |  | X |  |
| HALLMARK_PROTEIN_SECRETION |  |  |  |  |  |  |  |  |  |
| HALLMARK_REACTIVE_OXIGEN_SPECIES_PATHWAY |  |  |  |  |  |  |  | X |  |
| HALLMARK_SPERMATOGENESIS |  |  |  |  |  |  |  |  |  |
| HALLMARK_TGF_BETA_SIGNALING |  |  |  |  |  |  |  |  |  |
| HALLMARK_TNFA_SIGNALING_VIA_NFKB | X | X | X | X | X | X |  | X | X |
| HALLMARK_UNFOLDED_PROTEIN_RESPONSE |  |  |  |  |  |  |  | X | X |
| HALLMARK_UV_RESPONSE_DN |  |  |  |  |  |  |  | X | X |
| HALLMARK_UV_RESPONSE_UP | X |  |  |  | X | X |  | X | X |
| HALLMARK_WNT_BETA_CATENIN_SIGNALING |  |  |  |  |  |  |  | X |  |
| HALLMARK_XENOBIOTIC_METABOLISM | X |  |  | X |  | X | X | X | X |

**Table S8: Commonly altered pathways across cell lines panel.** Significantly enriched pathways from RNA-Seq data analysis are shown. X marks cell lines in which pathway was significantly up- or down-regulated.

**Supplementary Tables S9 and S10 included as excel files**

**Table S9: Transcription factor binding motifs enriched in less accessible chromatin regions.** Chromatin regions identified as less accessible after BRM011 treatment in THP-1 were queried for known transcription binding motifs. Transcription factor consensus sequence is shown together with significance score.

**Table S10: Transcription factor binding motifs enriched in less accessible chromatin regions.** Chromatin regions identified as more accessible after BRM011 treatment in THP-1 were queried for known transcription binding motifs. Transcription factor consensus sequence is shown together with significance score.
